# A Logic-incorporated Gene Regulatory Network Deciphers Principles in Cell Fate Decisions

**DOI:** 10.1101/2023.04.21.537440

**Authors:** Gang Xue, Xiaoyi Zhang, Wanqi Li, Lu Zhang, Zongxu Zhang, Xiaolin Zhou, Di Zhang, Lei Zhang, Zhiyuan Li

**Affiliations:** Peking-Tsinghua Center for Life Sciences, Academy for Advanced Interdisciplinary Studies, Peking University, Beijing, 100871, China; Center for Quantitative Biology, Academy for Advanced Interdisciplinary Studies, Peking University, Beijing, 100871, China; Beijing International Center for Mathematical Research, Center for Machine Learning Research, Peking University, Beijing 100871, China

**Keywords:** Gene Regulatory Network / Cell Fate Decision / Gene Regulatory Logic / Driving Force / Gene Expression Noise

## Abstract

Organisms utilize gene regulatory networks (GRNs) to make fate decisions, but the regulatory mechanisms of transcription factors (TFs) in GRNs are exceedingly intricate. A longstanding question in this field is how these tangled interactions synergistically contribute to decision-making procedures. To comprehensively understand the role of regulatory logic in cell fate decisions, we constructed a logic-incorporated GRN model and examined its behavior under two distinct driving forces (noise-driven and signal-driven). Under the noise-driven mode, we distilled the relationship among fate bias, regulatory logic, and noise profile. Under the signal-driven mode, we bridged regulatory logic and progression-accuracy trade-off, and uncovered distinctive trajectories of reprogramming influenced by logic motifs. In differentiation, we characterized a special logic-dependent priming stage by the solution landscape. Finally, we applied our findings to decipher three biological instances: hematopoiesis, embryogenesis, and trans-differentiation. Orthogonal to the classical analysis of expression profile, we harnessed noise patterns to construct the GRN corresponding to fate transition. Our work presents a generalizable framework for top-down fate-decision studies and a practical approach to the taxonomy of cell fate decisions.

## Introduction

Waddington’s epigenetic landscape is a fundamental and profound conceptualization of cell fate decisions [1]. Over decades, this insightful metaphor has facilitated researchers to distill a myriad of models regarding cell fate decisions [2–9]. While introducing various quantitative models and dissecting diverse fate-decision processes, researchers have further elaborated the Waddington landscape [10–15]. An outstanding question is whether the landscape is static or not, i.e., whether cell fate decisions are driven by noise or signal [14, 16, 17]. On one hand, some perspectives hold that cells reside in a stationary landscape, where decisions are made by switching through discrete valleys [18, 19], as a result of gene expression noise [20, 21]; termed as “noise-driven”, **Fig1.A**). Meanwhile, some researchers argued that the epigenetic landscape is dynamic during fate decisions. That is, the distortion of the landscape orchestrates fate transitions [7, 22, 23] and is driven by extrinsic signals (termed as “signal-driven”, **Fig1.B**).

**Figure 1.**
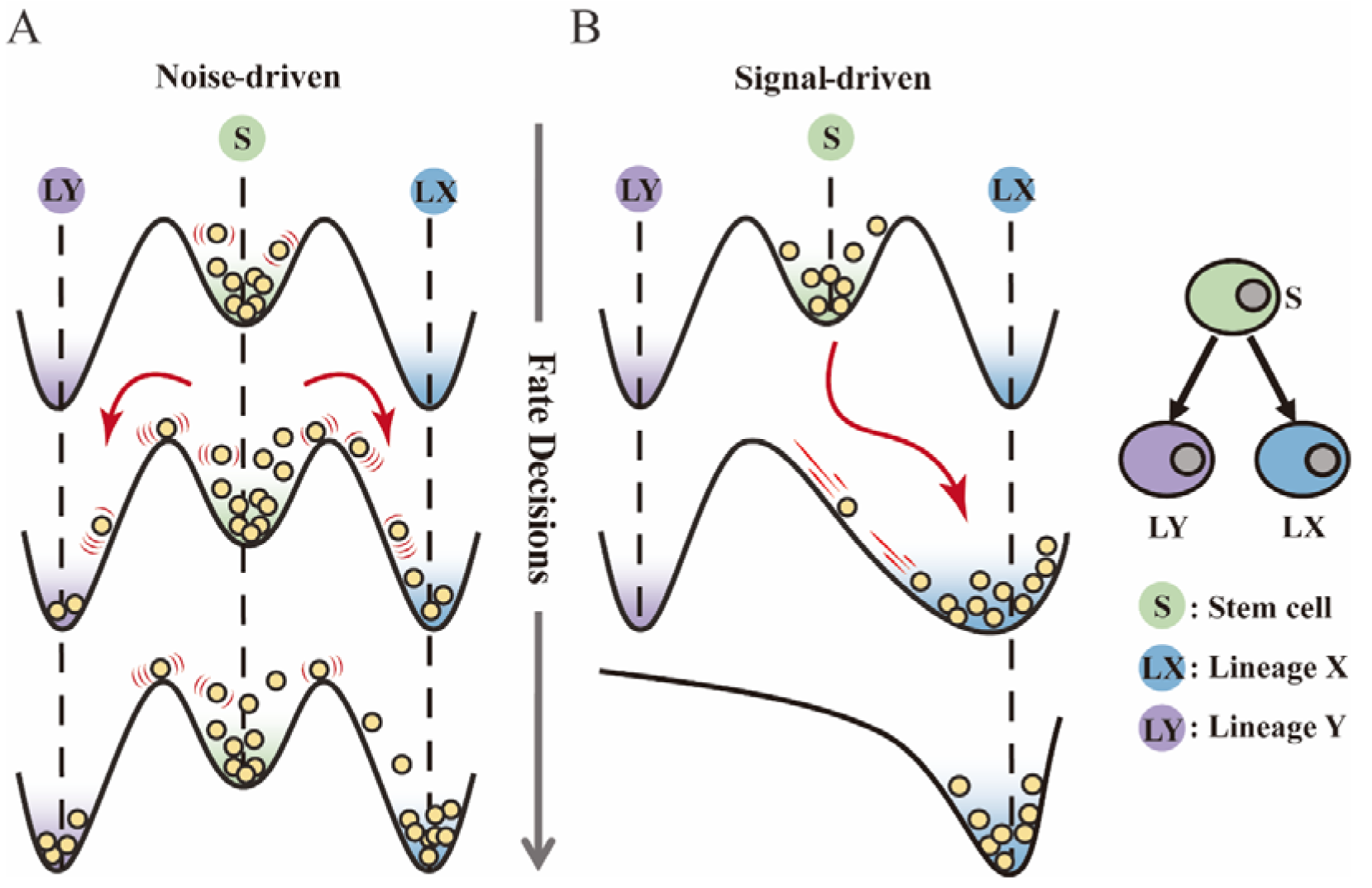
Schematic representation of cell fate decisions driven by noise (A) and signal (B) from a view of epigenetic landscape. (A-B) Valleys represent stable attractors. Cells (yellow balls) in stem cell fate (denoted as “S”, green well in landscape) differentiate into downstream fates, lineage X (denoted as “LX”, blue well) and lineage Y (denoted as “LY”, purple well). These abbreviations were used for following Figure 2-7.

Under the noise-driven mode, the bias of cell fate decisions largely depends on the spontaneous heterogeneity of gene expressions in the cell population [24, 25]. Consequently, the initial cellular state predominantly impacts the direction of the fate decision. Chang et al. [20] uncovered that hematopoietic stem cell (HSC) population possesses intrinsic and robust heterogeneity of *Scal-1* expression. Notably, populations with discrete expression levels of *Scal-1* confer different propensities for downstream lineage commitment. Considering the signal-driven mode, cell fates are tightly steered by extrinsic signals (e.g., cytokines, chemical molecules, mechanical strength and genetic operations) that reshape the landscape (**Fig1.B**). In this circumstance, the impact of the initial state on fate decisions is relevantly inconsequential. Additionally, due to the accessibility of signal manipulation, the signal-driven mode has been widely utilized for cell fate engineering [26, 27], leading to in-vitro induction systems centered on induced pluripotent stem cells (iPSCs) for obtaining desired cell types [28]. Recently, researchers reported a “fate-decision abduction” of erythroid-to-myeloid trans-differentiation induced by various type of cancer, which facilitates tumor escape from the individual’s immune system [29]. Collectively, driving forces couple the foundational and crucial features of fate decisions, serving as an essential basis for further decoding fate decisions and interpreting the development of organisms [16]. By examining the two driving modes, we can gain a better understanding and characterization of cell fate decisions, including in-vivo cell differentiation, oncogenesis, and in- vitro reprogramming systems.

Nevertheless, the driving forces that underlie fate decisions remain largely elusive. The intricate nature of gene regulatory networks (GRNs) presents a challenge in deciphering driving modes. It has been generally acknowledged that corresponding core GRNs orchestrate cell fate decisions [30, 31], where the lineage-specifying transcription factors (TFs) interact to implement fate-decision procedures. Furthermore, researchers transferred specific TFs into donor cells to reconfigure the intracellular GRNs for acquiring cell types of interest [32, 33]. Although some studies suggested that perturbation of a single TF is sufficient to transform certain cell fates [28], large number of TFs are inevitably involved in most differentiation/reprogramming processes [34]. In particular to orchestrate decisions among multiple cell fates, it is necessary for TFs to regulate target genes cooperatively [35]. As crucial determinants of cell fates, TFs function via binding to cis-regulatory elements (CREs, e.g., promoter and enhancer). CREs of a single gene in metazoans can simultaneously accommodate numerous TFs [36, 37]. While experimental protocols have been developed to assess TF binding and one-to-one up- or down-regulatory relationship, it is more challenging to quantify these combinatorial regulations. For instance, given two factors activate and inhibit the same target gene, respectively, does the target gene turn on or off when both factors are present in its CREs? Computational approaches in systems biology can be utilized to tackle complex networks [38–42]. A concise GRN model typically entails the following two elements. The element 1 is the topology. Much research efforts have been devoted to investigating network topologies on cellular behaviors, e.g., toggle switch [31, 43], and feed-forward loop [44–46]. In particular, the Cross-Inhibition with Self-activation (CIS) network is one of the most studied two-node GRNs in cell fate decisions [47, 48], with examples found in *GATA1*-*PU.1* and *FLI1*-*KLF1* in hematopoiesis [49, 50], *NANOG*-*GATA6* and *OCT4*-*CDX2* in gastrulation [16, 51], and *Sir2* and *HAP* in yeast aging [23]. In this topology, two lineage-specifying factors inhibit each other while active themselves. For example, in the well-known *GATA1*-*PU.1* circuit, *GATA1* directs fate of megakaryocyte-erythroid progenitor (MEP), and *PU.1* (also known as *SPI1*) specifies the fate of granulocyte-monocyte progenitor (GMP) [47]. Namely, the antagonism of two TFs implicates two cell fates in competition with each other.

The another element is the logic for regulatory functions [52, 53]. Exemplified by the CIS network, each node (e.g., X and Y) receives the activation by itself and inhibition by the counterpart. Hence there is naturally the logic function between these two inputs. Given the logic function is AND, in the context of biological mechanism of regulation by TFs, X gene expresses only when X itself is present in X’s CREs but Y is not. Researchers observed in the *E. coli* lac operon system that changes in one single base can shift the regulatory logic significantly, suggesting that logic functions of GRNs can be adapted on the demand of specific function in organisms [40, 54, 55]. Additionally, considering that the combination and cooperativity of TFs are of great significance in development [56–58], theoretical investigation of the logic underlying GRNs should be concerned in cell fate decisions. However, despite the existence of large number of mathematical models on fate decisions, the role played by the regulatory logic in cell fate decisions is still obscure. Some theoretical studies put emphasis on specific biological instances, adopting logic functions that best fit the observations derived from experiments [7, 39, 59]. As a result, the models incorporated different regulatory logic received limited attention. Other research assigned logic to large-scale multi-node GRNs, confining the interpretation of the role of logic [41, 60]. Collectively, the bridge between the logic of nodes in GRNs and cell fate decisions has not yet been elucidated systematically and adequately. Current research already encompassed a wealth of cell fate decisions: embryogenesis [61–63], lineage commitment [50, 64, 65], oncogenesis [66–68], in-vitro reprogramming [69–71], and large-scale perturbations [28, 72–74]. Analogous to the effort on taxonomy of cell types and tumors [75], how cell fate decisions can be classified and distilled to the common properties is a challenge for further exploring systematically and application on fate engineering [76, 77].

In this work, we integrated the fate-decision modes (noise-driven/signal-driven) and the classical logic operations (AND/OR) underlying GRNs in a continuous model. Based on our model, we investigated the impact of distinct logic operations on the nature of fate decisions with driving modes in consideration. Additionally, we extracted the difference in properties between the two driving modes. We unearthed that in the noise-driven stem cells, regulatory logic results in the opposite bias of fate decisions. We further distilled the relationship among noise profiles, logic motifs, and fate-decision bias, showing that knowledge of two of these allow inference of the third heuristically. Under the signal-driven mode, we identified two basic patterns of cell fate decisions: progression and accuracy. Moreover, based on our findings *in silico*, we characterized cell fate decisions in hematopoiesis and embryogenesis and unveiled their decision modes and logic motifs underlying GRNs. Ultimately, we applied our framework to a reprogramming system. We deciphered the driving force of this trans-differentiation, and utilized noise patterns for nominating key regulators. We underscored that clustering of gene noise patterns is an informative approach to investigate high-throughput dataset. Together, we underlined regulatory logic is of the significance in cell fate decisions. Our work presents a generalizable framework for classifying cell fate decisions and blueprint for circuit design in synthetic biology.

## Result

### Section 1: Mathematical model of the CIS network with logic motifs

Binary tree-like cell fate decisions are prevalent in biological systems [51, 75, 78, 79], orchestrated by a series of the CIS networks. Accordingly, we developed our ordinary differential equations (ODEs) model based on this paradigmatic and representative topology (Eq1, Eq2; see **Methods** for details).

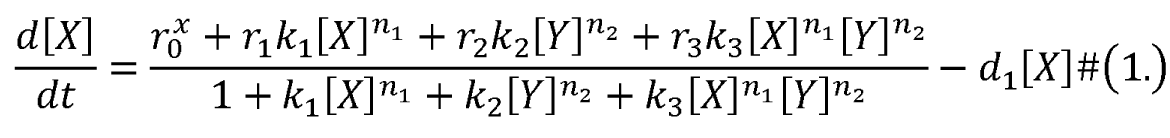

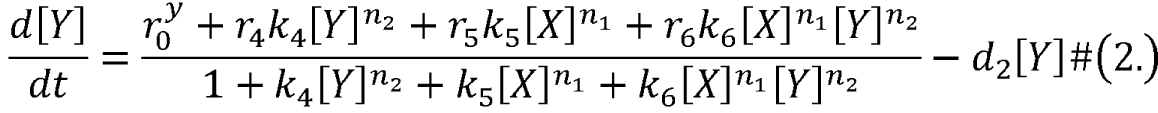

X and Y are TFs in the CIS network. *n_1_* and *n_2_*are the coefficients of molecular cooperation. *k_1_*-*k_3_* in Eq1 and *k_4_-k_6_* in Ep2 represent the relative probabilities for possible configurations of binding of TFs and CREs. (**Fig2.A**). *d_1_* and *d_2_* are degradation rates of X and Y, respectively. Here, we considered a total of four CRE’s configurations as shown in Figure 2A (i.e., TFs bind to the corresponding CREs or not, 2^2^=4). Accordingly, depending on the transcription rates (i.e., *r_0_^x^*, *r_1_*, *r_2_*, *r_3_* in Eq1, similarly in Eq2) of each configuration, we can model the dynamics of TFs in the Shea-Ackers formalism [80, 81].

**Figure 2.**
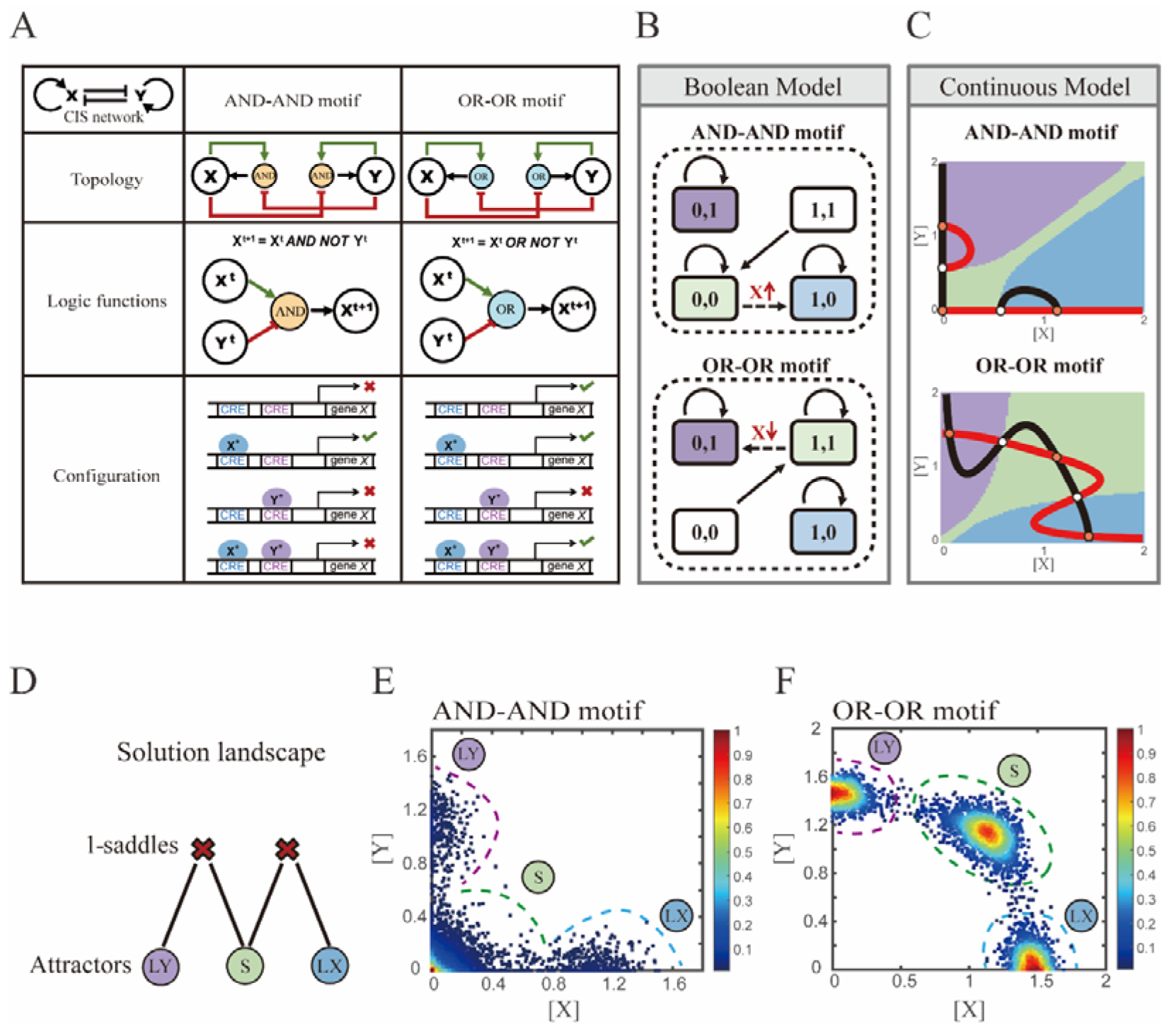
Models of the Cross-Inhibition with Self-activation (CIS) network incorporated logic motifs. (A) A table listing the topologies with logic nodes, logic functions and Cis-Regulatory Elements (CRE) configurations in the CIS network incorporated AND-AND and OR-OR logic (denoted as AND-AND motif and OR-OR motif). X and Y are lineage-specifying transcription factors (TF). X^t+1^ indicates the value of X at the next time step. X*, Y* represent activated forms of X and Y, respectively. The true or false signs denote whether gene *X* can be transcribed, respectively. These annotations were used for the following Figure 3-7. (B) State spaces of the AND-AND (top panel) and OR-OR (bottom panel) motifs in Boolean models. Updated rules of Boolean models are stated in Figure 2. Rectangles indicate cell states. Green, blue, purple represent S, LX, and LY, respectively. Solid arrows indicate transitions between states under corresponding Boolean models. Dotted arrows indicate forced transition imposed by external perturbations. (C) State spaces of the AND-AND (top panel) and OR-OR (bottom panel) motifs in ODE models. Dark and red lines represent nullclines of 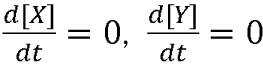, respectively. Stable steady states (SSS) are denoted as orange dots. Unstable Steady States (USSs) are denoted as white dots. Each axis represents the concentration of each transcription factor, which units are arbitrary. Blue, green and purple areas in state spaces indicate attractor basins representing LX, S and LY, respectively. Color of each point in state space was assigned by the attractors they finally enter according to the deterministic models (Eq1, Eq2). These annotations were used for the following Figure 3-7. (D) The solution landscape both for the AND-AND and OR-OR motifs. The crimson X-cross sign denotes the first-order saddle node. Blue, green, and purple circles indicate attractors. These annotations were used for the following Figure 3-7. (E-F) Simulation result of stochastic differential equation models of the AND-AND (E) and OR-OR (F) motifs. Other than adding a white noise, parameters were identical with those in (C). Initial values were set to the attractor representing S fate in Figure 2C top panel (E) and Figure 2C bottom panel (F). Noise levels of *X* (σ*_x_*) and *Y* (σ*_y_*) are both set to 0.14 in the AND-AND motif (E), and 0.1 in the OR-OR motif (F). Stochastic simulation was preformed 3500 times, with each final state recorded as a dot on the plot. Color of heatmap corresponds to the density of points. Unit of concentration is arbitrary.

Thus, the distinct logic operations (AND/OR) of two inputs (e.g., activation by X itself and inhibition by Y) can be further implemented by assigning corresponding profile of transcription rates in four configurations (**Fig2.A**). From the perspective of molecular biology, the regulatory logics embody the complicated nature of TF regulation that TFs function in a context-dependent manner. Considering the CIS network, when X and Y bind respective CREs concurrently, whether the expression of target gene is turned on or off depends on the different regulatory logics (specifically, off in the AND logic and on in the OR logic; **Fig2.A**). Notably, instead of exploring the different logics of one certain gene [44], we focus on different combinations of regulatory logics due to dynamics in cell fate decisions is generally orchestrated by GRN with multiple TFs.

Benchmarking the Boolean models with different logic motifs (**Fig2.B**; see **Methods**), we reproduced the geometry of the attractor basin in the continuous models resembling those represented by corresponding Boolean models (**Fig2.C**; see **Methods**). Under double AND and double OR motifs (termed as AND-AND motif and OR-OR motif, respectively), there are typically three stable steady states (SSSs) in the state spaces (**Fig2.C**): two attractors near the axes representing the fate of lineage X (denoted as LX, *X*^high^*Y*^low^) and the fate of lineage Y (denoted as LY, *X*^low^*Y*^high^), and the attractor in the center of the state space representing stem cell fate (denoted as S).

Evidently, the stem cell states exhibit different expression patterns between the two logic motifs. Stem cells in the AND-AND motif do not express *X* nor *Y* (**Fig2.B** top panel; express in low level in **Fig2.C** top panel), while in the OR-OR motif, stem cells express both lineage- specifying TFs (**Fig2.B** bottom panel; express in high level in **Fig2.C** bottom panel). The difference in the status of S attractors relates to the co-expression level of lineage-specifying TFs in stem cells in real biological systems [11, 50]. Intuitively, from the view of Boolean model, stem cell state in the AND-AND motif ([0,0] state) needs to switch on lineage-specifying TFs to transit to downstream fates (**Fig2.B** top panel). Whereas in the OR-OR motif, fate transitions are subject to the switch-off of TF expression (**Fig2.B** bottom panel). Furthermore, we introduced the solution landscape method. Solution landscape is a pathway map consisting of all stationary points and their connections, which can describe different cell states and transfer paths of them [82–84]. From the perspective of the solution landscape, two logic motifs possess akin geometric topologies in their steady-state adjacencies (**Fig2.D**): when there are three fates coexisting in the state space, S attractor resides in the middle of LX and LY as the possible pivot for fate transitions (**Fig2.D**). To investigate noise, we developed models with stochastic forms (see **Methods**). Simulations display the primary distribution of cell populations, corresponding to SSSs in deterministic models (**Fig2.E-F**).

### Section 2: Two logic motifs exhibit opposite bias of fate decisions under the noise-driven mode

We first investigated the difference between the AND-AND and OR-OR motifs under the noise- driven mode. Here, we assigned the stem cell state as the starting point in simulation. In biological systems, it is unlikely that the noise level of different genes is kept perfectly the same. Asymmetry of the noise levels was thus introduced. First, we set the noise level of TF *X* higher than that of *Y* (σ*_x_* =0.18, σ*_y_* = 0.12). Under this asymmetric noise, we observed that stem cells shifted toward LX in the AND-AND motif (**Fig3.A-B**), but toward LY in the OR-OR motif (**Fig3.C-D**). From the perspective of the state space, such properties intuitively originate from the distinctive status of stem cell attractors in two logic motifs (**Fig2.B-C**). In the AND-AND motif, the stem cell state resides at the origin of coordinates. Thus, with increasing *X*’s noise level, the stem cell population crossed the boundary between S and LX basins with a rising probability (**Fig2.C** top panel). Consequently, the fate decision of the stem cell population manifests a bias toward LX. Likewise, in the OR-OR motif, the stem cell population has a higher probability of entering LY basin following an increase in *X*’s noise (**Fig2.C** bottom panel). Next, we simulated multiple sets of noise levels for *X* and *Y*. We quantified distribution of cell types, which was determined by the basin in which the final state of each round of stochastic simulation ended up. We observed that stem cell population displays almost opposite differentiation preference under identical noise levels but distinct logic motifs (**Fig3.E**). Conversely, when two distinct logic motifs exhibiting the same fate-decision bias, cell populations need to employ opposite noise patterns (**Fig3.E-F**). Collectively, if two of the three (noise profiles, logic motif, fate-decision bias) are accessible, the last is inferential.

**Figure 3.**
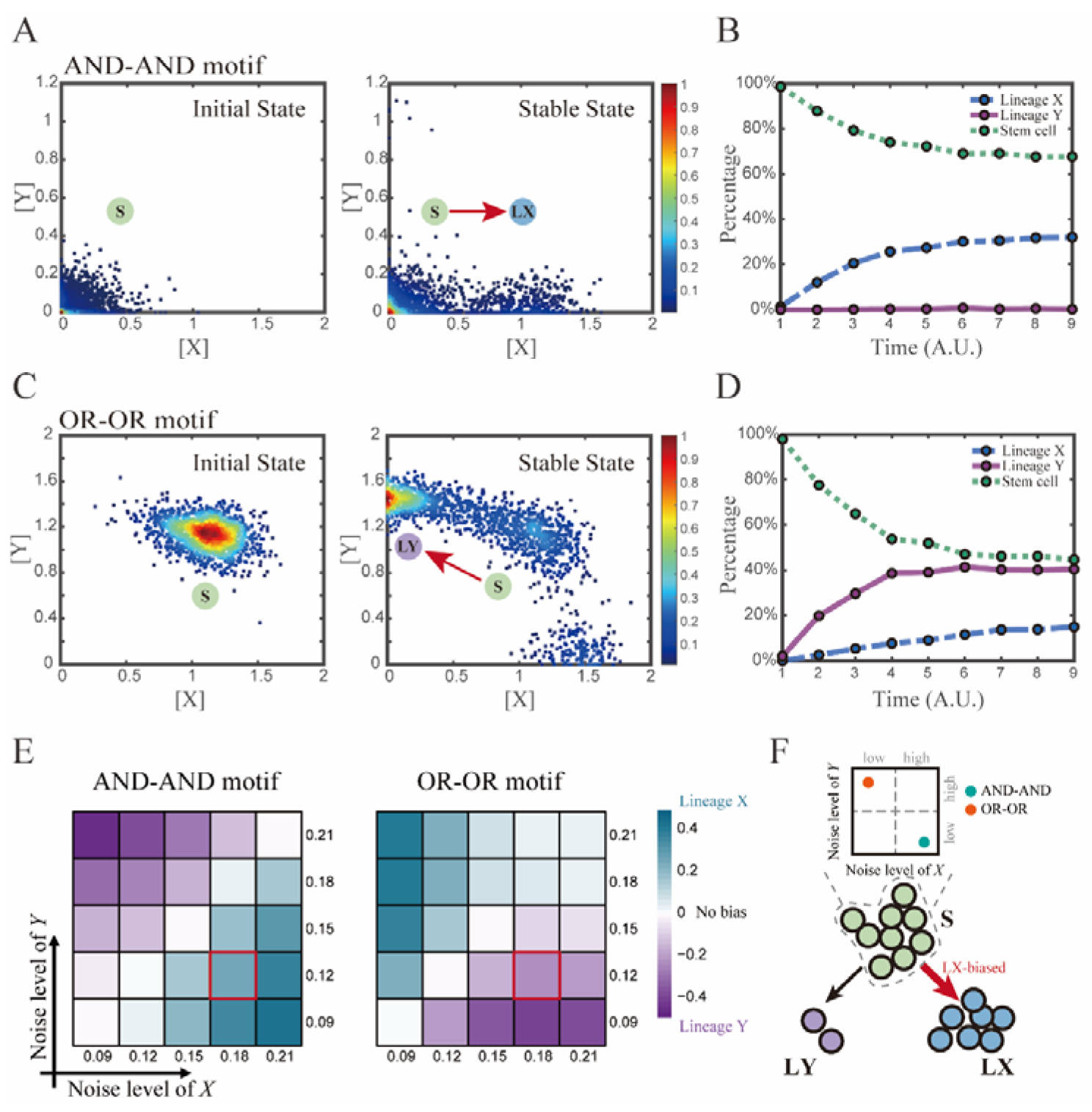
Two logic motifs exhibit opposite bias of fate decisions under the noise-driven mode. (A and C) Stochastic simulation in both the AND-AND and OR-OR motifs. σ*_x_* is set to 0.18, and σ_y_ is 0.12. In both (A) and (C), initial values were identical with attractors of stem cell fate in Figure 2C (SSSs in green attractor basins). Simulation was preformed 1500 times, with each initial (A left and C left) and final (A right and C right) states recorded as a dot on the plot. (B and D) Time courses of the percentage of cells in different fates in stochastic simulation, under the AND-AND motif (B) and OR-OR motif (D). Fates of cells were assigned by their final states according to the basins of the deterministic models in Figure 2C. Unit of time is arbitrary. (E) Heatmaps showing the bias of cell fate decisions under different noise levels of *X* and *Y*. Color of heatmap indicates the extent of bias. Here, bias 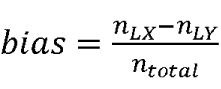. *n_LX_*, *n_LY_* represent number of LX, LY, respectively. *n_total_* represents the total number of cells (n_total_ = 1500). The method of assigning fate to cells is identical with Figure 3B and 3D. The red marked cells correspond to the noise conditions simulated in (A) and (C). (F) Schematic illustration in that stem cell populations possessing the same bias of fate decisions need to have opposite noise patterns, according to whether they are in the AND-AND or OR-OR motif. The red and bold arrow indicates the bias of fate decisions.

Next, we wondered whether noise could act as a driving force for reprogramming (e.g., from LY to S). We assigned LY state as the starting cell type in simulation. Apparently, in the AND- AND motif, transition of LY to S can be realized by increasing noise level of TF *Y* (**Fig**S1.A). Meanwhile in the OR-OR motif, it is the increased noise level of *X* that can drive the transition from LY to S (**Fig**S1.B), which is also intuitive by viewing the basin geometry of the state space (**Fig2.C**). These observations suggested that under the noise-driven mode, experimental reprogramming strategy need to take consideration of the regulatory logic (e.g., in reprogramming of LY to S, perturb the high expression TF of LY in the AND-AND motif, while in the OR-OR motif, perturb the low expression TF).

### Section 3: Two logic motifs decide oppositely between differentiation and maintenance under the signal-driven mode

In addition to noise, cell fate decisions can also be driven by signals, e.g., GM-CSF in hematopoiesis [85], CHIR99021 in chemically induced reprogramming [86]. The change conducted by signals corresponds to the distortions of the cell fate landscape. To simulate the signal-driven mode, we focused on the effect of parameters in the mathematical models on the system’s dynamical properties. To simulate models feasibly and orthogonally, we added parameters *u* (*u_x_*in Eq3, *u_y_* in Eq4) to Eq1-2:

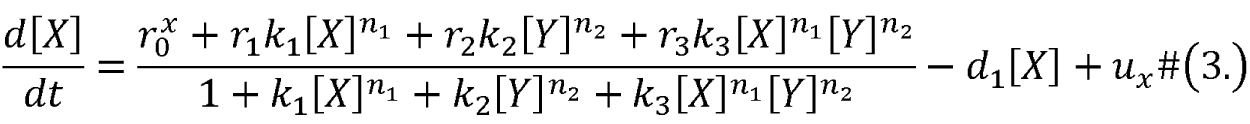

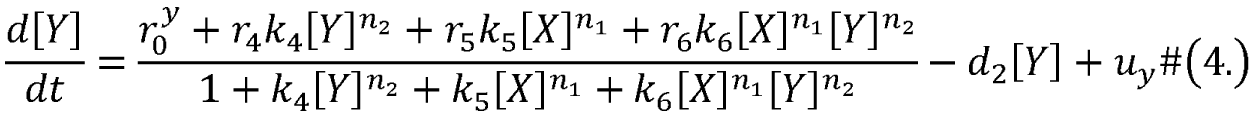

The increase of *u* represents an elevation in the basal expression level of lineage-specifying TFs, reflecting an induction signal from the extracellular environment. From an experimental standpoint, this signal can be the induction of small molecules or overexpression by gene manipulations, such as the transfection of cells with expression vectors containing specific genes.

We first explored the impact on the system when the two induction parameters are changed symmetrically (*u*=*u_x_*=*u_y_*). As the increase of *u*, the number of SSS in the AND-AND system decreases from three to two, where S attractor evaporates after a subcritical pitchfork bifurcation (**Fig4.A**, **Fig**S2.A). Whereas in the OR-OR motif, after the increase of *u*, LX and LY attractors disappear with saddle-node bifurcations respectively. Only the SSS representing stem cell fate is retained in the state space (**Fig4.B**). We then portrayed all the topology of the steady-state adjacency that accompanied the increase of *u,* from the perspective of the solution landscape. In the AND-AND motif, the attractor basin of LX and LY started to adjoin and occupied the vanishing S attractor basin together (**Fig4.C-D**). Accordingly, the stem cells cannot maintain themselves and decided to differentiate into either one of the lineages. Moreover, if the cell population possesses the same noise levels in both *X* and *Y*, then the fate decisions are unbiased (**Fig**S2.B).

**Figure 4.**
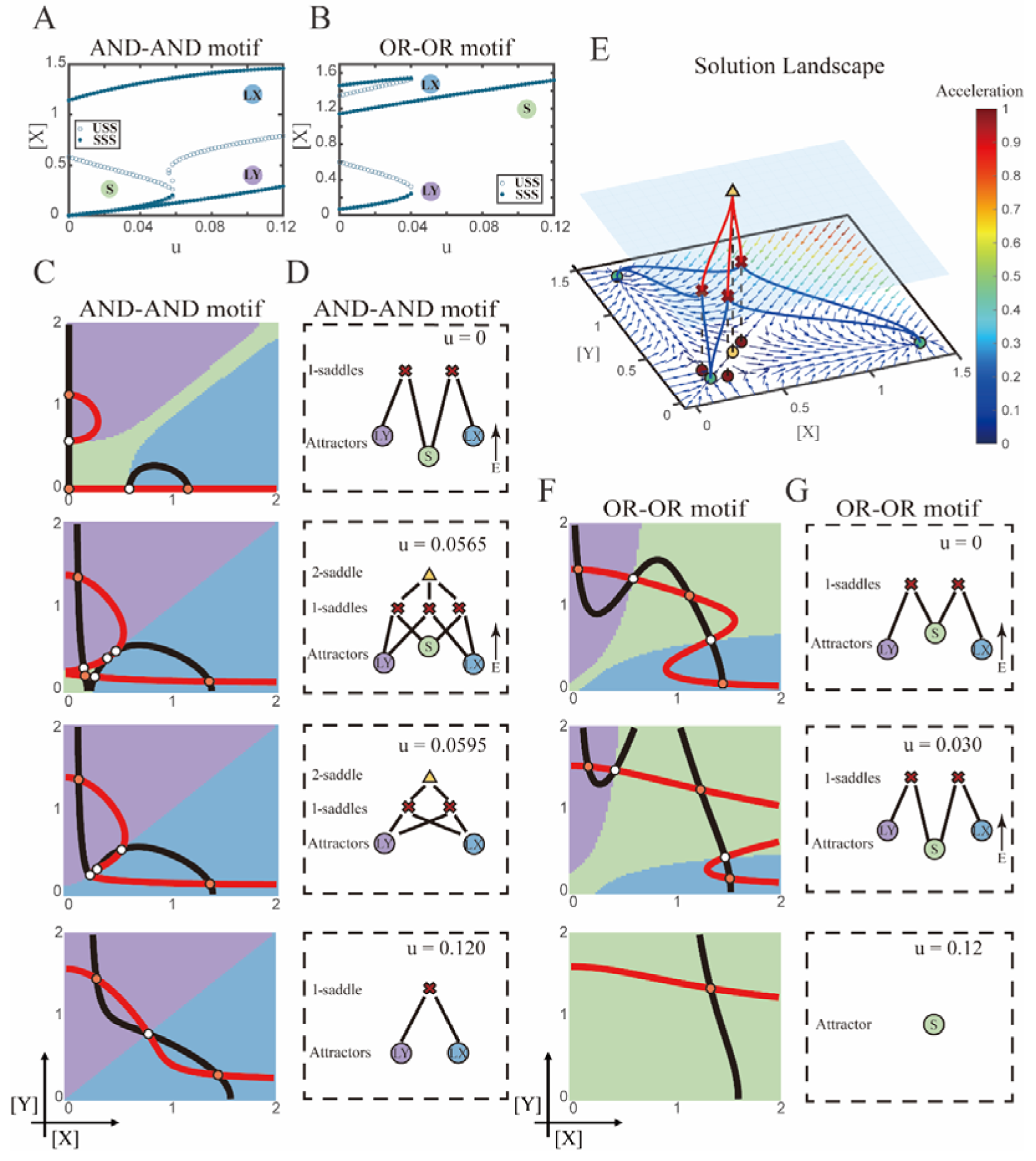
Two logic motifs decide oppositely between differentiation and maintenance under the signal- driven mode. (A-B) Bifurcation diagrams for the AND-AND motif (A) and OR-OR motif (B) driven by parameter *u* (*u* = *u_x_* = *u_y_*) in the CIS model. SSSs and USSs are denoted as solid dots and hollow dots, respectively. (C and F) Changes in the state spaces for the AND-AND motif (C) and OR-OR motif (F) with increasing parameter *u*, from top to down. (D and G) Changes in the solution landscape with increasing of *u*, in company with these in (C and F). The crimson X-cross sign and yellow triangle denote first-order and second-order saddle nodes, respectively. Relative energy is quantified by the geometric minimum action method [90], see **Methods**. (E) The solution landscape with parameter *u* = 0.0565 for the AND-AND motif from a view of three dimensions. It describes a hierarchical structure of the steady states. From top to bottom, it represents 2-saddle (yellow triangle), 1-saddles (crimson X-cross sign), and the attractors (green dot). The layer of 1-saddles is represented by a blue translucent plane, and the bottom layer is the flow field diagram. The connections from 2-saddle to 1-saddles are represented by red lines, and the connection from 1-saddles to the attractors are represented by blue lines. In the flow field diagram, the direction and color of the arrows correspond to the direction and size of the flow at that location. The corresponding positions of 2-saddle and 1-saddles in the flow field are marked with yellow and red dots, respectively, with black dashed lines indicating the corresponding relationship.

Notably, in the AND-AND motif we observed a brief intermediated stage before S attractor disappears, where all three fates are directly interconnected (**Fig4.C** 2^nd^ panel and D 2^nd^ panel, **Fig.4E**). To manifest the generality, we globally screened 6,213 groups of parameter sets under the AND-AND motif, and this logic-dependent intermediated stage can be observed for 82.7% of them (see **Methods;** Table S1), indicating little dependence on particular parameter setting (1.8% in the OR-OR motif). Unlike the indirect attractor adjacency structure mediated by S attractor (**Fig2.D**), the solution landscape with fully-connected structure facilitates transitions between any two pairs of fates. Furthermore, this transitory fully-connected stage locates between the fate- undetermined stage (**Fig4.C** top panel) and fate-determined stage (**Fig4.C** 3^rd^ panel), comparable to the initiation (or activation) stage before the lineage commitment in experimental observations [87–89]. Therefore, we suspected that the robust fully-connected stage in the AND-AND motif may correspond to a specific period in cell fate decisions.

From the standpoint of reprogramming of differentiated cells back into progenitors, in the AND-AND motif, differentiated cells are more capable of maintaining their own fates during the symmetrical increase of the induction signals on both lineages (**Fig4.A** and C). Whereas in the OR-OR motif, the attractor basin of LX or LY is progressively occupied by the stem cell fate as *u_x_*and *u_y_* increase together (**Fig4.F-G**). In this scenario, the downstream fates are eventually reversed back to the undifferentiated state (**Fig4.B** and F). Namely, reprogramming engaged in the OR-OR motif can be accomplished by bi-directional induction of downstream antagonistic fates. In sum, we found that under symmetrical signal induction, the behavior of stem cells is subject to core GRN’s logic motifs. In the AND-AND motif, stem cells prefer to differentiate, while under the OR-OR motif, the stem cell population inclines to maintain its undifferentiated state.

### Section 4: The trade-off between progression and accuracy of cell fate decisions under the signal-driven mode

According to experimental observations, the majority of fate decisions exhibit lineage preference, also known as “symmetry breaking of fate decisions” [14, 64, 65, 91]. Take the lineage choices in hematopoiesis as an example, Some HSCs prefer myeloid over lymphoid [64, 92]. This fate-decision bias also further shifts along with aging and infection [88, 93]. In studying this preference in fate decisions, we broke the symmetry in the signal-driven models, by solely increasing *u_x_* while keeping *u_y_* =0 (**Fig5.A**). First, it is apparent that the fate decision will significantly steer toward LX along with the increase of *u_x_*, regardless of the logic motifs. Ultimately the state spaces contain only LX attractor when *u_x_* is sufficiently high (**Fig5.B-C**, **Fig**S3.A). However, the changes in the state space and the solution landscape follow different routes for two logic motifs. In the AND-AND motif, S attractor basin disappears at first, leaving a state space with two differentiated fates (**Fig5.D-E**). Then the basin of LY attractor shrinks and finally disappears (**Fig5.D-E**). Whereas in the OR-OR motif, LY attractor disappears first. Then S attractor, with an enlarged basin, shares the state space with LX attractor. Finally, S attractor basin abruptly disappears by a saddle-node bifurcation (**Fig5.F-G**).

**Figure 5.**
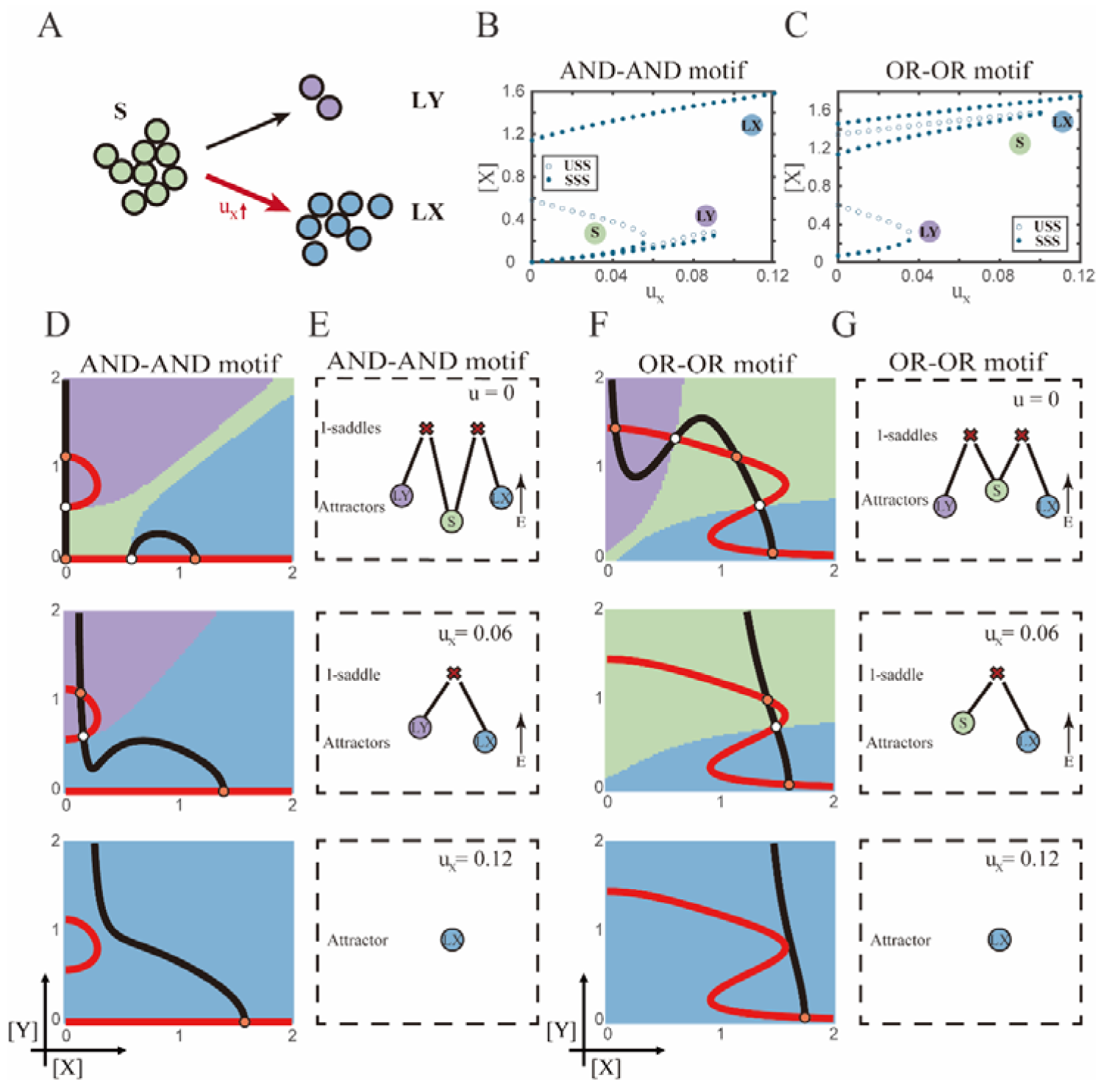
The progression-accuracy trade-off in cell fate decisions. (A) Schematic illustration of S-to-LX cell fate decisions with *X*-inducing signals. The red and bold arrow indicates the direction of fate decisions. (B-C) Bifurcation diagrams for the AND-AND motif (B) and OR-OR motif (C) driven by parameter *u_x_*. (D and F) Changes in the state spaces for the AND-AND motif (D) and OR-OR motif (F) with increasing values of *u_x_*, from top to down. (E and G) Changes in the solution landscape with increasing of *u_x_*, in company with these in (D and F).

The distinct sequences of attractor basin disappearance as *u_x_* increasing can be viewed as a trade-off between progression and accuracy. In the AND-AND motif, the attractor basin of LX and LY adjoins (**Fig5.D** middle panel) when S attractor disappears due to the first saddle-node bifurcation (**Fig5.B**). Notwithstanding the bias of differentiation toward LX, the initial population still possesses the possibility of transiting into LY (**Fig**S3.B). That is, in the AND-AND motif, as the increase of induction signal *u_x_*, the “gate” for the stem cell renewal is closed first. Stem cells are immediately compelled to make fate decisions toward either LX or LY, with a bias toward LX but a nonignorable probability of entering LY. Albeit the accuracy of differentiation is therefore compromised, the overall progression of differentiation is ensured (i.e., all stem cells have to make the fate decisions downward. This causes the pool of stem cells to be exhausted rapidly). Whereas in the OR-OR motif, the antagonistic fate, LY, disappears first (**Fig5.C**). The attractor basin of S and LX are adjacent in the state space (**Fig5.F**). In this case, the orientation of the fate decisions is generally unambiguous since the stem cell population can only shift to LX, ensuring the accuracy of differentiation. Next, to check if the observed sequences of basin disappearance are artifacts of specific parameter choice, we randomly sampled parameter sets to check the sequence of attractor changes in their state spaces (6,207 groups of the AND-AND motifs and 6,634 groups of the OR- OR motifs; Table S1). We found that 96% AND-AND motifs and 70% OR-OR motifs exhibit the same sequence of attractor vanishment mentioned above (**Fig**S3.C-D; see **Methods**). These results

of the global screen demonstrated that the sequence of attractor vanishment is robust to parameter settings. In sum, we proposed that logic motifs couple the trade-off between progression and accuracy as a general phenomenon in the signal-driven asymmetrical fate decisions (**Fig**S3.E).

Next, we examined the trans-differentiation from LY into LX by increasing *u_x_*. In the AND- AND motif, with the induction of *X*, LY directly transited into LX as the stem cell state disappears before LY (**Fig5.D**, **Fig**S4.A). Intriguingly, for the OR-OR motif under the same induction, LY population first returned to the S state and then flows into LX (**Fig5.F**, **Fig**S4.B). Namely, different logic motifs conduct distinct trajectories in response to identical induction in reprogramming. The AND-AND motif renders a one-step transition between downstream fates (**Fig**S4.C). While in the OR-OR motif, it is a two-step transition mediated by the stem cell state (**Fig**S4.D). This phenomenon suggests the observation that cells may be reprogrammed to distinct cell types depending on the induction dose [86] is more realizable in the OR-OR motif. Integrated with the foregoing symmetrical induction, we recapitulated that in the OR-OR motif, the bi-directional induction or a unilateral induction from a counterpart (e.g., solely induced *Y* to realize reprogramming of LX to S) confer downstream cell fates to return to the undifferentiated state (**Fig4.F**, **Fig5.F**). Whereas in the AND-AND motif, it is substantially more difficult to achieve de- differentiation. This observation may explain why some cell types are not feasible to reprogram [94].

### Section 5: The CIS network performs differently during hematopoiesis and embryogenesis

In prior sections, we systematically investigated two logic motifs under the noise- and signal- driven modes *in silico*. With various combinations of logic motifs and driving forces, features about fate-decision behaviors were characterized by computational models. Next, we questioned whether observations in computation can be mapped into real biological systems. And how to discern different logic motifs and driving modes is a prerequisite for answering this question.

To end this, we first evaluated the performance of different models, specifically in simulating the process of stem cells differentiating towards LX (**Fig6.A**). Under four models with different combinations of driving modes and logic motifs (**Fig**S5.A-B), we assessed the expression level and expression variance (defined as the coefficient of variation) of TFs *X* and *Y* among the cell population over time in stochastic simulation. We observed that, under the same logic motifs, different driving modes change in the patterns of expression variance rather than expression levels (**Fig6.B-C**, **Fig**S5.C-D). Overall, under the noise-driven differentiation from S to LX, the variance of expression exhibits a continuous and monotonic trend (**Fig**S5.D) for both logic motifs. For different logic motifs, in the AND-AND motif, the expression variance of *X* (highly expressed in LX) declines (**Fig**S5.D top panel). Whereas in the OR-OR motif, it is the expression variance of *Y* (low expressed in LX) displays a rising trend (**Fig**S5.D bottom panel). Nevertheless, under the signal-driven mode, the expression variance increases and then decreases, exhibiting a non- monotonic transition due to signal-induced bifurcation. During S to LX differentiation, comparable to the noise-driven mode, it is the expression variance of TF *X* in the AND-AND motif and TF *Y* in the OR-OR motif display a nonmonotonic pattern.

**Figure 6.**
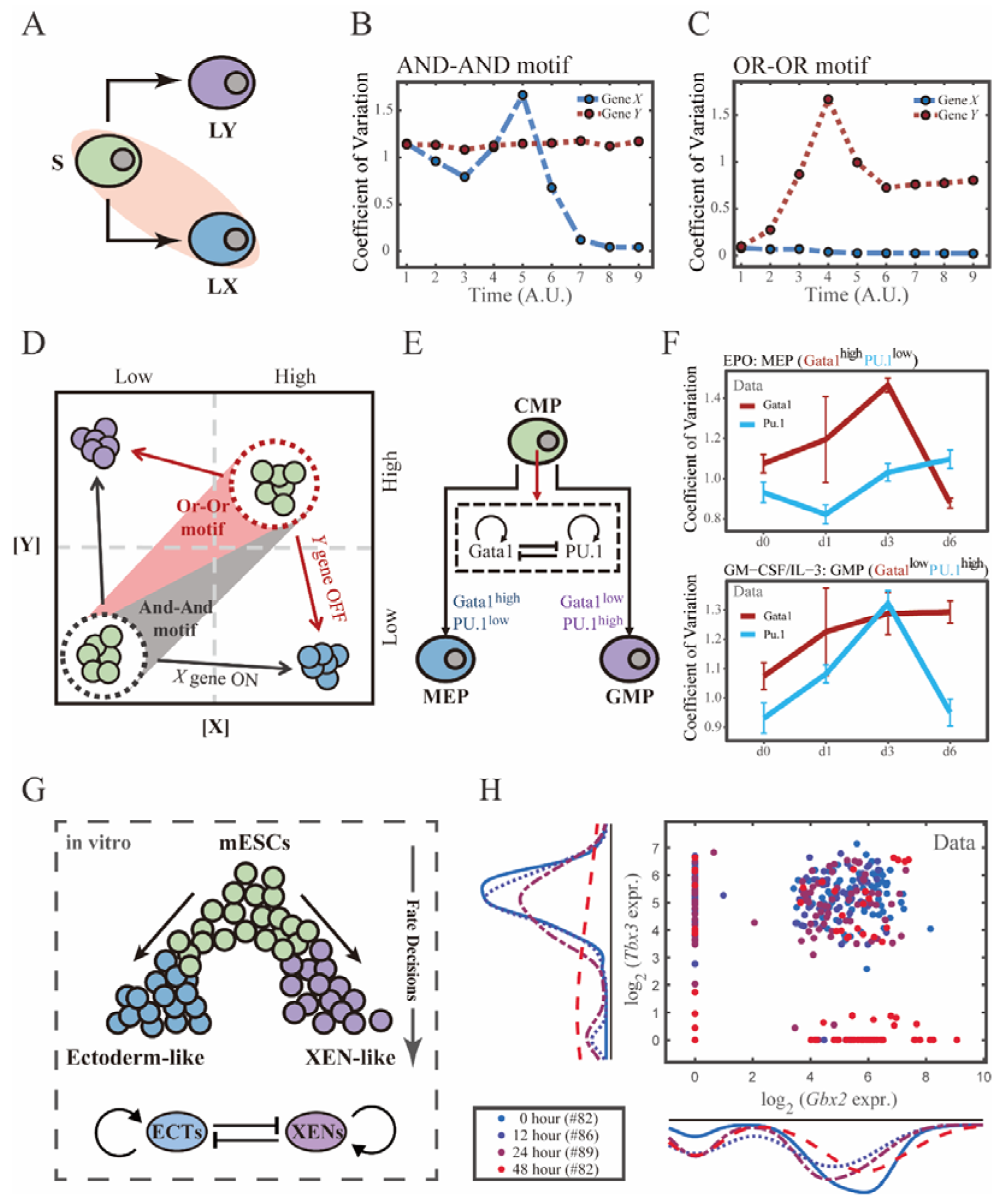
The CIS network performs differently during hematopoiesis and embryogenesis. (A) Schematic illustration of S differentiating into LX. We took fate transition labeled in light pink shade as an example in following simulation. (B) Time courses on the coefficient of variation in expression levels of *X* and *Y* genes *in silico* during differentiation towards LX (*u_x_*switches from 0 to 0.08 from time point 1 to 9) in the AND-AND motif. Initial values were set to the attractors of stem cell fate in Figure 2C top panel (SSS in green attractor basin). σ*_x_* and σ*_y_* are both set to 0.07. Stochastic simulation was preformed 1000 times for each pseudo-time point. Unit of time is arbitrary. (C) Time courses on the coefficient of variation in expression levels of *X* and *Y* genes *in silico* during differentiation towards LX (*u_x_*switches from 0 to 0.24 from time point 1 to 9) in the OR-OR motif. Initial values were set to the attractors of stem cell fate in Figure 2C bottom panel (SSS in green attractor basin). σ*_x_* and σ*_y_* are both set to 0.05. Stochastic simulation was preformed 1000 times for each pseudo-time point. Unit of time is arbitrary. (D) Schematic illustration of distinctive cell fate decision patterns under the AND-AND and OR-OR motifs in the state space. Dark and red gradients represent the extent of “AND-AND” and “OR-OR” in the actual regulatory network, respectively. Each axis represents expression levels of the lineage-specifying TFs. Blue, green, and purple circles indicate the cell fates of LX, S, and LY, respectively. (E) Schematic illustration of *Gata1-PU.1* circuit that dominates the primary fate decisions in hematopoiesis (CMP: Common myeloid progenitor; MEP: megakaryocyte-erythroid progenitor; GMP: Granulocyte-monocyte progenitor). (F) Measured coefficient of variation of expression levels of *Gata1* and *PU.1* changing over time during differentiation from CMPs to MEPs and GMPs. Expression levels were quantified via single-cell RT-qPCR [85]. Error bars on points represent standard deviation (SD). For details of data processing, see **Methods.** (G) Schematic illustration of the differentiation from mESCs in induction system [95]. (H) Measured expression levels of *Gbx2* and *Tbx3* among cells in embryogenesis quantified via single-cell SMART-seq2 [95]. For details of data processing, see **Methods.**

Such disparities between logic motifs originate from the location of S attractor (**Fig6.D**, **Fig2.C**). Although the target cell types are the same (LX), the AND-AND motif requests the expression of the TF *X* to be turned on, while the OR-OR motif requests the TF *Y* to be turned off. These key fate-transition genes, namely TF *X* in the AND-AND motif and TF *Y* in the OR-OR motif, both exhibit a sharp increase of variation in response to saddle-node bifurcation driven by *u_x_* induction (1^st^ saddle node in **Fig5.B**; 2^nd^ saddle node in **Fig5.C**). Overall, these computational results suggest that we may be able to distinguish the two driving modes according to the expression variance over time series, then logic motifs can be correspondingly assigned by the expression level of the genes in the target cell types. For instance, if the expression variance of the *X* gene exhibits a nonmonotonic pattern and *X* is highly expressed in target cell types, then this cell fate decision can be assigned as the signal-driven fate decision in an AND-AND-like motif.

To support our findings with real-world correspondence, we first focused on the differentiation of CMPs in hematopoiesis (**Fig6.E**). It is acknowledged that the transcriptional regulation of *Gata1*-*PU.1* circuit dominates this cell fate decisions, which conforms to the CIS topology (**Fig6.E**). Mojtahedi et al. [85] stimulated murine multipotent hematopoietic precursor cell line EML with erythropoietin (EPO) or granulocyte macrophage colony-stimulating factor/interleukin 3 (GM-CSF/IL-3) to examine the commitment into an erythroid or a myeloid fate, respectively. Based their curated dataset, we found that the expression level of *Gata1* (highly expressed in MEPs) gradually increased during EPO induction (**Fig**S6.A), while the expression variance exhibits a nonmonotonic trend (**Fig6.F** top panel). Symmetrically, during GM-CSF/IL-3 induced differentiation toward GMPs, the expression level of *PU.1* (highly expressed in GMPs) gradually increased (**Fig**S6.B), while the expression variance also presents a nonmonotonic pattern (**Fig6.F** bottom panel). The trends shown in the dataset resembles the signal-driven mode with the AND-AND motif. In addition, we quantified the expression of *Gata1* and *PU.1* via the single molecule FISH dataset (**Fig**S6.C) [24]. They are at low levels in CMPs, corresponding to the expression patterns in the AND-AND motifs (**Fig2.E**, **Fig6.D**). Together, we suggested that the *Gata1*-*PU.1* circuit performs in an AND-AND-like manner, and this differentiation system [85] is under the signal-driven mode.

Another paradigmatic model of fate decision is the differentiation of embryonic stem cells (ESC). Semrau et al. [95] found that under the retinoic acid (RA) exposure system in vitro, mouse embryonic stem cells (mESCs) differentiated into two lineages: extraembryonic endoderm (XEN)- like and ectoderm-like. The investigators recapitulated that two clusters of TFs with the CIS topology determined this lineage specification (**Fig6.G**). We observed that the expression variance in most of these fate-decision TFs (16/22 73%) are gradually increasing during time, and 14% (3/22) of them exhibit nonmonotonic behavior (**Fig**S5.D), suggesting the process is more likely driven by noise (**Fig**S6.E). Furthermore, we focused on potential key regulators: *Gbx2* and *Tbx3*, the two likely targets of RA that are crucial for this fate decision [95]. The expression variances over time of these two TFs are consistently increasing (**Fig**S6.D). In addition, their initial expressions are at high level, in agreement with that of the OR-OR motif (**Fig6.D** and H). In short, we proposed that the mESCs differentiation system under RA exposure performs in an OR-OR- like manner, and its differentiation is under the noise-driven mode in this experimental setting.

### Section 6: The chemical-induced reprogramming of human erythroblasts (EBs) to induced megakaryocytes (iMKs) is the signal-driven fate decisions with an OR-OR-like motif

The foregoing cell fate decisions initiate from pluripotent cells in mice (mESCs, CMPs) corresponding to a typical "downhill" in Waddington’s metaphor. In 2006, Yamanaka et al. accomplished the reprogramming from mouse fibroblasts into iPSC state via the noted "OSKM" factors, representing "uphill" in Waddington’s metaphor [94, 96, 97]. Likewise, trans- differentiations from one lineage to another have been realized by overexpression or chemical inductions [26], whether they correspond to direct “trespassing” of the ridge or an “up-and-down” through the peak in Waddington landscape are still elusive. We then applied our models to reprogramming systems, with a primary focus on hematopoiesis. Qin et al. [98] recently achieved the direct chemical reprogramming of EBs to iMKs using a four-small-molecule cocktail (**Fig7.A**). Investigators presented that EBs underwent an induced bipotent precursor for erythrocytes and MKs (iPEMs) to finally desired iMKs. It is acknowledged that the *FLI1-KLF1* circuit with the CIS topology dominates this fate-decision process [47, 50]. To deduce the logic motif of the *FLI1- KLF1* circuit, we quantified the expression patterns of *FLI1* and *KLF1* based on published single- cell RNA-seq data [98]. We can observe the fate transition from the EB population (*FLI1*^low^, *KLF1*^high^) to the iMK population (*FLI1*^high^, *KLF1*^low^) (**Fig7.B**). According to their expression level, the cell populations can be primarily classified into three clusters. In addition, both *FLI1* and *KLF1* are highly expressed in the intermediate cell population suspected to be the progenitors of iMKs and EBs [98]. Namely, the pattern of expression level is concordant with the OR-OR motif in our framework (**Fig6.D**).

**Figure 7.**
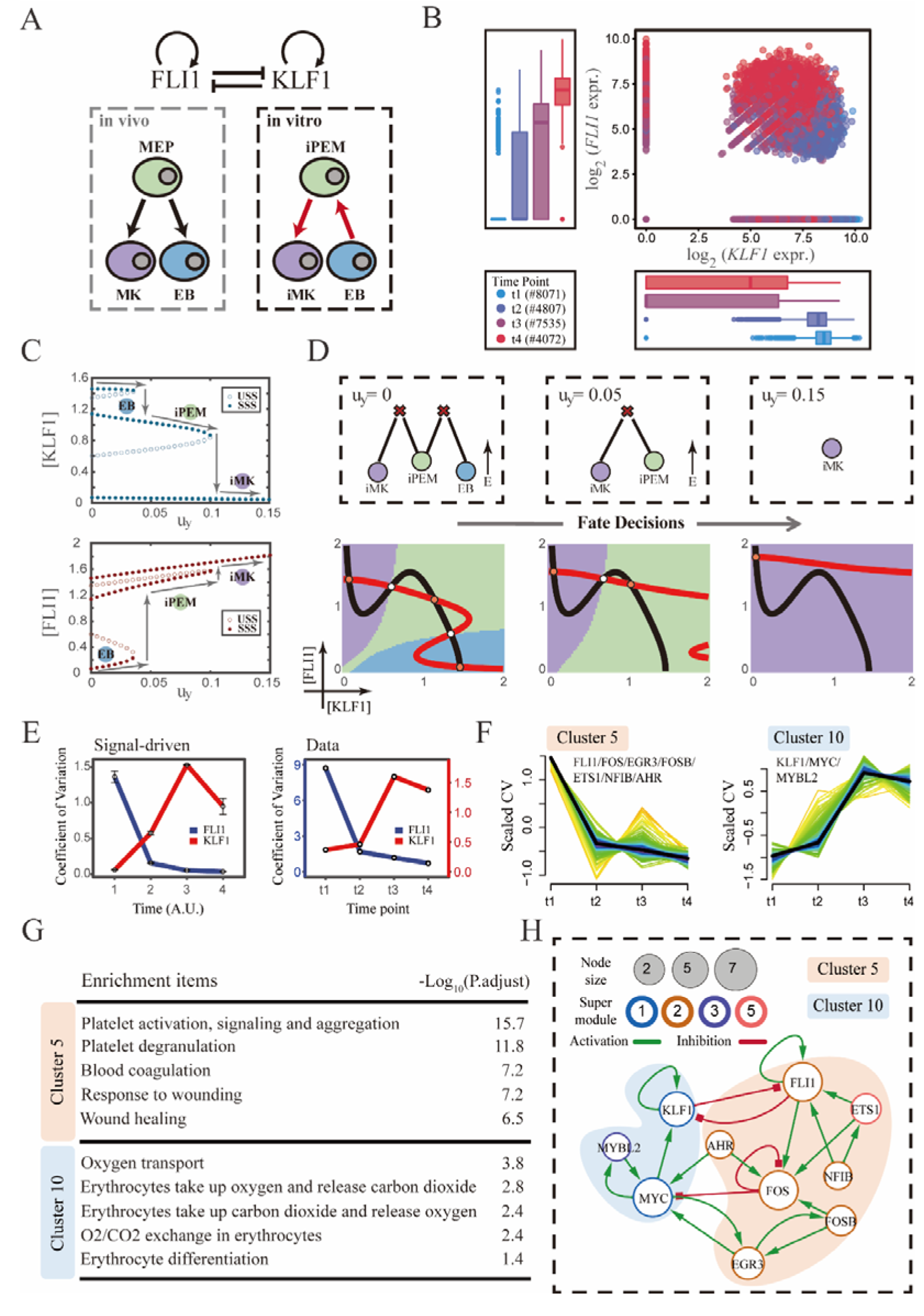
The chemical-induced reprogramming of human EB to iMK is the signal-driven fate decisions with an OR-OR-like motif. (A) Schematic illustration of the differentiation from MEPs in vivo and in vitro. Red arrows represent the route of reprogramming [98] (B) Measured expression levels of *KLF1* and *FLI1* in reprogramming quantified via single-cell 10X. For details of data processing, see **Methods.** (C) Bifurcation diagrams for the OR-OR motif driven by parameter *u_y_* in the CIS model. (D) Fate transition representing reprogramming of EB to iMK *in silico*. Top panel: changes in the solution landscape with increasing of parameter *u_y_*, from left to right; Bottom panel: changes in the state spaces for the OR- OR motif with increasing values of *u_y_*, in company with these in top panel. Unit of concentration is arbitrary. (E) Left panel: coefficient of variation of expression levels of *KLF1* and *FLI1* changes *in silico* over time under given parameter (*u_y_*= 0.11) in the OR-OR motif. Noise level of *KLF1* (σ*_x_*) and *FLI1* (σ_y_) are set to 0.087. Initial values were identical with LX attractor in Figure 2C bottom panel (SSS in blue attractor basin). Stochastic simulation was preformed 1000 times per round for each time point. We totally preformed 3 round simulations. Error bars on points represent SD; Right panel: measured coefficient of variation of expression levels of *KLF1 and FLI1* changing over time in the processes from EBs to iMKs. Unit of time is arbitrary. (F) Identification of distinct temporal patterns of expression variance by fuzzy c-means clustering. The x axis represents four time points, while the y axis represents scaled CV (coefficient of variation) in each time point. Dark trend lines in the middle indicate the average of scaled CV over genes in cluster. (G) Enriched major Gene Ontology terms for cluster 5 and 10. (H) Regulatory network of TFs in cluster 5 and 10. Circle size indicates the sum of in-degree and out-degree. Node colors indicate different Supermodules (adapted from [98]). Green and red edges indicate activation and inhibition, respectively. The light blue and light pink shades denote genes in cluster 5 and 10, respectively.

Next, to investigate the driving force of this reprogramming system, we simulated the fate transition from EB (corresponding to LX, blue) to iMK (LY, purple) under both driving modes. Under the noise-driven mode, we assumed that the reprogramming system facilitated the noise levels of both TFs. For simplicity, the starting cell population (EBs) was assigned symmetrical high noise levels. While under the signal-driven mode, we assumed that the four-small-molecule cocktail upregulated the expression of *FLI1* (highly expressed in iMKs). Then, we simulated the transition from EBs to iMKs by lifting the basal expression level of *FLI1*, corresponding to parameter *u_y_*in model (**Fig7.C**). The bifurcation diagrams indicate that the signal-driven fate transition is mediated by the iPEM state (**Fig7.C**). In particular, overexpression of *FLI1* renders sequential saddle-node bifurcations. Thus, EBs are converted to iPEMs before steering toward the terminal iMK state (Fig7.C and D), which is consistent with the experiment’s findings.

Furthermore, we next assessed the dynamic behaviors in models of this system under different driven modes with the OR-OR logic. As discussed before, there is no discernible difference in the expression levels between the two driving modes. Overall, the common tendency is an up-regulation in *KLF1* and a down-regulation in *FLI1* (**Fig**S7.A). We then quantified the expression variances during this fate transition by the model. Under the noise-driven mode, expression variances of *FLI1* and *KLF1* would gradually decrease and increase, respectively, until stabilizing (**Fig**S7.B). Under the signal-driven mode, the expression variance of *FLI1* would first decline and then remain nearly constant, while *KLF1* would exhibit a nonmonotonic pattern (**Fig7.E** left panel). From the view of modeling, the nonmonotonic pattern presented by *KLF1* originates from the rapid shut-off during the transition from iPEM state to iMK state (2^nd^ saddle node in **Fig7.C**).

Accordingly, we next quantified the expression variances in the real dataset over time. Impressively, the pattern emerging from the data accommodates the hypothesis of the signal- driven mode (**Fig7.E**). Altogether, we proposed that this reprogramming system [98] is the signal- driven process underlying the OR-OR-like motif. Moreover, the high expression level of *FLI1* induced by small molecules is the key driving force of the fate transition, as suggested by the properties of the OR-OR motif. We underlined that it is a classical two-step fate-decision process mediated by the upstream progenitor state, which is in agreement with the phenomenon articulated by Qin et al. [98].

We then searched for genes with similar patterns of expression variance to those of *KLF1* and *FLI1*, with a hypothesis that genes possessing comparable expression variance patterns, especially TFs, may synergistically perform fate-decision related functions. Thus, we applied the fuzzy c- means algorithm [99] to cluster genes based on their expression variances, rather than expression levels. In total, we observed 12 distinct clusters of temporal patterns (**Fig**S7.C; Table S2). We focused primarily on clusters 5 and 10, where *KLF1* and *FLI1* are found respectively (**Fig7.F**). To testify our hypothesis, we conducted an enrichment analysis using the gene set in cluster 5 and 10. Functions associated with specific cell types are significantly enriched (**Fig7.G**). In particular, cluster 5 with *FLI1* is largely related to platelet-related functions, whereas cluster 10 with *KLF1* is largely related to the energy metabolism of blood cells. Furthermore, we filtered out 15 TFs in cluster 5 and 10 (11 in cluster 5; 4 in cluster 10). Next, by harnessing the TF interaction database (see **Methods**; Table S3), we collected 21 regulons associated with 10 TFs to construct TF regulatory Network (**Fig7.H**). Intriguingly, in the original article, genes were classified into six “Supermodules” according to the patterns of expression levels. Genes in cluster 5 are located in Supermodules 1 and 3, representing a decreasing tendency for expression level. Meanwhile, genes in cluster 10 are distributed in Supermodule 2 and 5, and their expression levels raise from low to high (Figure 6.D from [98]). Of note, most of TFs filtered by expression variance patterns appear in the GRN constructed in the original article (8/10, 80%; Figure 6.G from [98]). As a result, we underscored that *EGR3* and *ETS1*, which did not appear in the GRN of the original article, have been suggested to play important roles in the trans-differentiation. In addition, we observed that *MYC* and *FOS* possess the largest connectivity in our 10-node GRN, suggesting that these two TFs as hubs are essential regulators of this reprogramming system.

Together, in mapping our framework to the real reprogramming system, we assigned the cell fate decisions of EBs to iMKs to the signal-driven mode incorporated the OR-OR-like motif, by comparing the expression and expression variance patterns measured from real dataset with pseudo-data produced by our models in a “top-down” fashion. According to the model, the reprogramming is primarily driven by induced up-regulation of FLI1. Additionally, from the view of expression variance, we recapitulated a concise 10-node GRN and identified some TF nodes not previously recognized as major regulators, like *EGR3* and *ETS1*.

## Discussion

Comprehending the driving forces behind cell fate decisions is crucial for both fundamental scientific research and biomedical engineering. Recent advances in data collection and statistical methods have greatly enhanced our understanding of the mechanisms that regulate cell fate.

However, despite the widespread use of the Waddington landscape as a metaphor in experiments, there has been little examination into whether the fate decision observed in a particular experiment corresponds to a stochastic shift from one attractor to another on the landscape (i.e., noise-driven) or an overall distortion of the landscape (i.e., signal-driven). The application of appropriate computational models in systems biology can aid in uncovering the underlying mechanisms [100, 101]. One of the most representative work is that Huang et al. [59] modeled the bifurcation in hematopoiesis to reveal the lineage commitment quantitatively. Compared to simply modularizing activation or inhibition effect by employing Hill function in previous work, our models reconsidered the multiple regulations from the level of TF-CRE binding.

Our computational investigations have emphasized the importance of the combinatorial logic in connecting gene expression patterns with the driving forces underlying cell fate decisions.

Utilizing a representative network topology known as the CIS, our analysis demonstrated how both driving forces and regulatory logic jointly shape expression patterns during fate transitions. In turn, mean and variance in gene expression patterns can reveal logic motifs and driving forces. Our analytical framework promotes the interpretability of fate decisions and can be employed to speculate on the driving factors of fate decisions using a "top-down" approach, thereby providing a reference for investigating the causality of fate decisions and experimental validation.

The roles of noise as a possible driver of fate transitions are intriguing. By our models, the relationship among noise configuration, logic motifs, and fate-decision bias has been unveiled, and we noticed the opposite fate bias for the AND-AND and OR-OR motifs. Conversely, on the demand of the same bias, the progenitors with different logic motifs tend to employ a different profile of noise level (**Fig3.E-F**). Therefore, we suggested that under the noise-driven mode, the logic motif works like a “broker” to shape the fate preferences. Based on the assumption that the preference of fate decisions is the result of evolution and adaptation, we posited that if an organism has a functional demand for a specific bias, the noise profile will be iterated via logic motifs. In turn, changes in noise levels will be mediated by logic motifs to shape the differentiation bias. One intuitive example is the significant shift in differentiation preferences of HSCs over aging [102–104]. It has been reported that aging HSCs show different level of DNA methylation and epigenetic histone modifications from young HSCs [92, 105, 106]. How changes in epigenetic level shape the noise profile of the cell population and further affect the shift of fate- decision preference is a fascinating question. We underscored that the dissection of logic motifs underlying associated GRNs is a prerequisite for answering that question.

With the ever-expanded of single-cell sequencing data, characterizing genes by their mean expressions does not make the most of high-throughput datasets. As an intrinsic characteristic in central dogma, expression variance has been utilized to locate the “critical transitions” in complex networks [107, 108], e.g., identify the critical transitions in diseases like lymphoma [109]. Instead of differentially expressed genes, Rosales-Alvarez et al. [110] harnessed differentially noisy genes to characterize the functional heterogeneity in HSC aging. Our work presents that the patterns of expression variance can also be used to indicate driving forces and key regulators during the fate- decision processes. Nevertheless, compared to traditional gene expression, the interpretation of expression variance patterns is generally not intuitively accessible [111]. Additionally, extra a priori knowledge is needed to filter out the cluster of interest. To this end, our framework enables researchers to locate functional clusters via mathematic model based on appropriate hypothesis. Notably, if the genes that constituting the CIS network are not specified, we can conversely leverage the patterns of temporal expression variance to nominate key regulators in a model- guided manner. Collectively, our framework provides a mechanistic explanation for expression variance patterns and qualitatively characterize key expression variance patterns to locate core regulators of fate decisions without reliance on a priori knowledge.

Comparing to tuning noise, altering signals is a more accessible approach for experimentally manipulating cell fates. When the basal expressions of two lineage-specifying genes grow symmetrically, we have shown that opposite trends of fate transitions occur under two logic motifs: In the AND-AND motif, it promotes differentiation, whereas the OR-OR motif stabilizes stem cell fates. This is reminiscent of the "seesaw" model where maintenance of stemness can be achieved by overexpression of antagonistic lineage-specifying genes [112]. Our model suggests that restoring stemness by inducing two antagonistic lineage-specifying genes is more likely under the OR-OR-like motif (**Fig**S2.C). This is in concert with our analysis that mESC differentiation system performs in an OR-OR-like manner. In addition, Mojtahedi et al. [85] found that, under simultaneous induction of two antagonistic fates (EPO and GM-CSF/IL-3), although differentiation was delayed, it eventually occurred. This observation is consistent with models in the AND-AND motif, and we suggested that the core regulatory circuits in hematopoiesis performs in an AND-AND-like manner. More experimental validations would be needed to validate this hypothesis that “the seesaw model prefers the OR-OR motif”. Conversely, insight from the "seesaw" model also provides a candidate approach to further testify logic motifs underlying GRNs in experiments. For instance, stem cells with an AND-AND-like circuit are expected to display differentiation rather not maintenance in response to bidirectional induction.

In quantifying the signal-driven landscape changes, the solution landscape enables intuitive interpretation even in high-dimension GRNs [82, 113, 114]. In this work, we used both the state space and the solution landscape, in order to relate them for further investigations involving more than two TFs. Interestingly, from the perspective of the solution landscape, we found a robust fully-connected stage in the AND-AND motif. We envisioned that this period corresponds to the priming stage of differentiation. Notably, this fully-connected stage was not found in the OR-OR motif, suggesting that the necessity for priming during differentiation may be subject to the logic motifs of core GRNs.

Actual cell fate decisions are seldom purely unbiased. Under the asymmetrical signal-driven mode, we summarized the progression-accuracy trade-off in cell fate decisions: If a large number of cells are ensured to differentiate, then concessions have to be made in the accuracy of differentiation, and vice versa (**Fig**S3.E). An intuitive example is the large-scale apoptosis occurs daily in hematopoiesis. Hence, maintaining homeostasis in vivo inevitably requests cells to respond rapidly to differentiation. A recent study reported that in response to systemic inflammation by polymicrobial sepsis, pool of CMPs is rapidly depleted to accelerate the production of downstream cell fates [115]. This is concordant with our result that *Gata1*-*PU.1* circuit in hematopoiesis performs in an AND-AND-like manner. On the other hand, in embryonic development like *C.elegans*, the accuracy of cell fate decisions is considerably emphasized. How this nature of differentiation has been adapted in evolution is an interesting question. Our work highlights that this property is associated with the logic motifs of GRNs, suggesting the emphasis on progression or accuracy may be embedded in the logic motifs of core GRNs.

We classified three examples of cell fate decisions based on patterns of expression and expression variance. In hematopoiesis, we took fate choice between erythroid and myeloid as a paradigm, and assigned it an AND-AND-like motif under the signal-driven mode. In embryogenesis, we suggested the fate decision in RA exposure system is an OR-OR-like motif under the noise-driven mode. In reprogramming, the chemical-induced trans-differentiation is the signal-driven fate decisions incorporated an OR-OR-like motif. For simplicity and intuitiveness, we devised our model with two symmetrical combinations of regulatory logic (AND-AND/OR- OR). Albeit there are merely four types of cell fate decisions in consideration, our framework enables to be generalized and expanded to accommodate multi-node GRNs and complex logic combinations. Plenty of studies zoomed in one particular fate-decision events. However, from the standpoint of systems biology, we underlined that classification of fate decisions is a vital step for further investigation, as is the case for the typing of cells and tumors. Theoretically, appropriate classification of fate-decision systems enables the enrichment of common properties. So, accumulated knowledge can be inherited to new fate-decision cases. Taking reprogramming as an example, Zhao [86] recapitulated five kinds of trajectories in chemical-induced reprogramming. We suggested that the reprogramming trajectory is coupled with the logic motifs. On one hand, it is possible to answer why a certain reprogramming system exhibit a particular trajectory. On the other hand, it is possible to postulate achievable reprogramming according to the logic motifs of core GRNs (e.g., the AND-AND motif is more likely to enable direct conversion; model 4 mentioned in [86]). Recently, synthetic biology has realized the insertion of the CIS network in mammalian cells [22]. One of the prerequisites for recapitulating the complex dynamics of fate transitions in synthetic biology is systematical understanding of the role of GRNs and driving forces in differentiation. And the logic motifs are the essential and indispensable elements in GRNs. Our work also provides a blueprint for designing logic motifs with particular functions. We are also interested in validating the conclusions drawn from our models in a synthetic biology system.

## limitation of this study

Although our framework enables the investigation of more logic motifs, we chose two classical and symmetrical logic combinations for our analysis. Future work should involve more logic gates like XOR and explore asymmetrical logic motifs like AND-OR. The gene expression datasets analyzed here are only available for a limited number of time points. Though they meet the need for discerning trends, it is evident that the application to the datasets with more time points will yield clearer and less ambiguous changing trends to support the conclusions of this paper more generally. Notwithstanding the fact that the CIS network is prevalent in fate-decision programs, there are other topologies of networks that serve important roles in the cell-state transitions, like feed-forward loop, etc. The framework should further incorporate diverse network motifs in the future. In addition, for simplicity and intuition, we here considered signals as uncoupled and additive effects in ODE models, due to feasible mapping in real biological systems, such as ectopic overexpression.

## Supplemental information

Supplemental information includes seven figures and three tables and can be found with this article online.

## Acknowledgements

We thank Y. Zhao for advice of model application on real biological process; D. Grün for insightful and generous feedback on noise interpretation; J. Zhang, Y. Li, and X. Pei for providing original raw-data processing R scripts; Z. Zhou for illustrations and useful discussion; and the entire Zhiyuan laboratory for support and advice.

## Funding

This work was supported by the National Key Research and Development Program of China (No. 2021YFF1200500, 2021YFA0910700), and National Natural Science Foundation of China (No.12225102, 12050002, and 12226316). It is also supported by grants from Peking-Tsinghua Center for Life Sciences.

## Author contributions

Conceptualization, G.X. and Z.L.; Methodology, G.X., Xiaoyi Z. and Z.Z.; Software, G.X. and Xiaoyi Z.; Formal Analysis, G.X.; Investigation, G.X.; Resources, G.X. and D.Z.; Data Curation, G.X. and W.L.; Writing – Original Draft, G.X.; Writing – Review & Editing, G.X., Xiaoyi Z., W.L., Lu Z., Z.Z., Xiaolin Z., Lei Z and Z.L.; Visualization, G.X., Xiaoyi Z. and Z.Z.; Supervision, Lei Z. and Z.L.; Project Administration, Lei Z. and Z.L.; Funding Acquisition, Lei Z. and Z.L.

## Declaration of interest

The authors declare no competing interests.

## Notes

### Competing Interest Statement

The authors have declared no competing interest.

### Summary of Updates

The following is a summary of the major changes: a) According to the recommendations from reviewers, we have made necessary adjustments to the use of terminologies to improve clarity of our work: i) we have substituted AND-AND/OR-OR for original expression of AA/OO; ii) we now employ "transitory fully-connected stage" instead of "temporal fully-connected stage"; iii) regarding the term "Pulse-like behavior", we now adopt the terms "monotonic transitions" and "nonmonotonic transitions" to underline the distinct temporal noise's patterns in cell fate decisions brought by two driving forces in a more contrastive way; b) We have added an additional section titled "limitation of this study" to the revised manuscript, and explicitly pointed to the potential limitations of our work including those mentioned by reviewers. In addition, we have further polished the manuscript including our computational models, decisions regarding parameter choices, and logic flows in each section; c) We modified the unclear figures, legends, and phrasing in the main text. In addition, we have included more extensive discussions regarding the potential impact of our work on the field, with appropriate citations.

## References

1. Waddington, C.H., The strategy of the genes: A discussion of some aspects of theoretical biology. 1957, London: Allen & Unwin.

2. MacArthur, B.D., The geometry of cell fate. Cell Syst, 2022. 13(1): p. 1–3.

3. Schiebinger, G., et al., Optimal-Transport Analysis of Single-Cell Gene Expression Identifies Developmental Trajectories in Reprogramming. Cell, 2019. 176(4): p. 928–943 e22.

4. Olsson, A., et al., Single-cell analysis of mixed-lineage states leading to a binary cell fate choice. Nature, 2016. 537(7622): p. 698-702.

5. Bhattacharya, S., Q. Zhang, and M.E. Andersen, A deterministic map of Waddington’s epigenetic landscape for cell fate specification. BMC Systems Biology, 2011. 5(1): p. 85.

6. Fisher, A.G. and M. Merkenschlager, Fresh powder on Waddington’s slopes. EMBO reports, 2010. 11(7): p. 490–492.

7. Hota, S.K., et al., Brahma safeguards canalization of cardiac mesoderm differentiation. Nature, 2022. 602(7895): p. 129-134.

8. Huang, S., The molecular and mathematical basis of Waddington’s epigenetic landscape: A framework for post-Darwinian biology? BioEssays, 2012. 34(2): p. 149–157.

9. Li, C.J., et al., MicroRNA governs bistable cell differentiation and lineage segregation via a noncanonical feedback. Molecular Systems Biology, 2021. 17(4).

10. Shakiba, N., et al., How can Waddington-like landscapes facilitate insights beyond developmental biology? Cell Syst, 2022. 13(1): p. 4–9.

11. Loeffler, D. and T. Schroeder, Understanding cell fate control by continuous single-cell quantification. Blood, 2019. 133(13): p. 1406–1414.

12. Coomer, M.A., L. Ham, and M.P.H. Stumpf, Noise distorts the epigenetic landscape and shapes cell-fate decisions. Cell Syst, 2022. 13(1): p. 83–102 e6.

13. Shi, J., et al., Energy landscape decomposition for cell differentiation with proliferation effect. National Science Review, 2022: p. nwac116.

14. Stanoev, A. and A. Koseska, Robust cell identity specifications through transitions in the collective state of growing developmental systems. Current Opinion in Systems Biology, 2022. 31.

15. Glauche, I. and C. Marr, Mechanistic models of blood cell fate decisions in the era of single-cell data. Current Opinion in Systems Biology, 2021. 28.

16. Simon, C.S., A.K. Hadjantonakis, and C. Schröter, Making lineage decisions with biological noise: Lessons from the early mouse embryo. WIREs Developmental Biology, 2018. 7(4).

17. Zernicka-Goetz, M. and S. Huang, Stochasticity versus determinism in development: a false dichotomy? Nat Rev Genet, 2010. 11(11): p. 743–744.

18. Lord, N.D., et al., Stochastic antagonism between two proteins governs a bacterial cell fate switch. Science, 2019. 366(6461): p. 116.

19. Desai, R.V., et al., A DNA-repair pathway can affect transcriptional noise to promote cell fate transitions. Science, 2021: p. eabc6506.

20. Chang, H.H., et al., Transcriptome-wide noise controls lineage choice in mammalian progenitor cells. Nature, 2008. 453(7194): p. 544-7.

21. Guillemin, A. and M.P.H. Stumpf, Noise and the molecular processes underlying cell fate decision-making. Phys Biol, 2021. 18(1): p. 011002.

22. Zhu, R., et al., Synthetic multistability in mammalian cells. Science, 2022. 375(6578): p. eabg9765.

23. Li, Y., et al., A programmable fate decision landscape underlies single-cell aging in yeast. Science, 2020. 369(6501): p. 325.

24. Wheat, J.C., et al., Single-molecule imaging of transcription dynamics in somatic stem cells. Nature, 2020. 583(7816): p. 431-436.

25. Kovary, K.M., et al., Expression variation and covariation impair analog and enable binary signaling control. Molecular Systems Biology, 2018. 14(5).

26. Xu, J., Y. Du, and H. Deng, Direct lineage reprogramming: strategies, mechanisms, and applications. Cell Stem Cell, 2015. 16(2): p. 119–34.

27. Del Vecchio, D., et al., A Blueprint for a Synthetic Genetic Feedback Controller to Reprogram Cell Fate. Cell Syst, 2017. 4(1): p. 109–120 e11.

28. Ng, A.H.M., et al., A comprehensive library of human transcription factors for cell fate engineering. Nat Biotechnol, 2021. 39(4): p. 510–519.

29. Long, H., et al., Tumor-induced erythroid precursor-differentiated myeloid cells mediate immunosuppression and curtail anti-PD-1/PD-L1 treatment efficacy. Cancer Cell, 2022. 40(6): p. 674–693.e7.

30. Qiu, X., et al., Mapping transcriptomic vector fields of single cells. Cell, 2022.

31. Graf, T. and T. Enver, Forcing cells to change lineages. Nature, 2009. 462(7273): p. 587-94.

32. Hammelman, J., et al., Ranking reprogramming factors for cell differentiation. Nat Methods, 2022. 19(7): p. 812–822.

33. Mazid, M.A., et al., Rolling back of human pluripotent stem cells to an 8-cell embryo-like stage. Nature, 2022.

34. Hersbach, B.A., et al., Probing cell identity hierarchies by fate titration and collision during direct reprogramming. Molecular Systems Biology, 2022. 18(9).

35. Trojanowski, J. and K. Rippe, Transcription factor binding and activity on chromatin. Current Opinion in Systems Biology, 2022. 31.

36. Lambert, S.A., et al., The Human Transcription Factors. Cell, 2018. 172(4): p. 650–665.

37. Reiter, F., S. Wienerroither, and A. Stark, Combinatorial function of transcription factors and cofactors. Curr Opin Genet Dev, 2017. 43: p. 73–81.

38. Macarthur, B.D., A. Ma’ayan, and I.R. Lemischka, Systems biology of stem cell fate and cellular reprogramming. Nat Rev Mol Cell Biol, 2009. 10(10): p. 672–81.

39. Krumsiek, J., et al., Hierarchical differentiation of myeloid progenitors is encoded in the transcription factor network. PLoS One, 2011. 6(8): p. e22649.

40. Collombet, S., et al., Logical modeling of lymphoid and myeloid cell specification and transdifferentiation. Proc Natl Acad Sci U S A, 2017. 114(23): p. 5792–5799.

41. Moignard, V., et al., Decoding the regulatory network of early blood development from single- cell gene expression measurements. Nat Biotechnol, 2015. 33(3): p. 269–276.

42. Kirouac, D.C., et al., Cell–cell interaction networks regulate blood stem and progenitor cell fate. Molecular Systems Biology, 2009. 5(1).

43. Davies, J., et al., An IRF1-IRF4 Toggle-Switch Controls Tolerogenic and Immunogenic Transcriptional Programming in Human Langerhans Cells. Front Immunol, 2021. 12: p. 665312.

44. Kittisopikul, M. and G.M. Suel, Biological role of noise encoded in a genetic network motif. Proc Natl Acad Sci U S A, 2010. 107(30): p. 13300–5.

45. Oliver Metzig, M., et al., *An incoherent feedforward loop interprets NF*κ*B/RelA dynamics to determine TNF*LJ*induced necroptosis decisions*. Molecular Systems Biology, 2020. 16(12).

46. Sciammas, R., et al., An incoherent regulatory network architecture that orchestrates B cell diversification in response to antigen signaling. Molecular Systems Biology, 2011. 7(1).

47. Orkin, S.H. and L.I. Zon, Hematopoiesis: an evolving paradigm for stem cell biology. Cell, 2008. 132(4): p. 631–44.

48. Kato, H. and K. Igarashi, To be red or white: lineage commitment and maintenance of the hematopoietic system by the "inner myeloid". Haematologica, 2019. 104(10): p. 1919–1927.

49. Hoppe, P.S., et al., Early myeloid lineage choice is not initiated by random PU.1 *to GATA1 protein ratios.* Nature, 2016. 535(7611): p. 299-302.

50. Palii, C.G., et al., Single-Cell Proteomics Reveal that Quantitative Changes in Co-expressed Lineage-Specific Transcription Factors Determine Cell Fate. Cell Stem Cell, 2019. 24(5): p. 812–820 e5.

51. Zhou, J.X. and S. Huang, Understanding gene circuits at cell-fate branch points for rational cell reprogramming. Trends Genet, 2011. 27(2): p. 55–62.

52. Ocone, A., et al., Reconstructing gene regulatory dynamics from high-dimensional single-cell snapshot data. Bioinformatics, 2015. 31(12): p. i89–i96.

53. Matsuura, S., et al., Synthetic RNA-based logic computation in mammalian cells. Nat Commun, 2018. 9(1).

54. Mayo, A.E., et al., Plasticity of the cis-regulatory input function of a gene. PLoS Biol, 2006. 4(4): p. e45.

55. Buchler, N.E., U. Gerland, and T. Hwa, On schemes of combinatorial transcription logic. Proc Natl Acad Sci U S A, 2003. 100(9): p. 5136.

56. Zaret, K.S., Pioneer Transcription Factors Initiating Gene Network Changes. Annu Rev Genet, 2020. 54: p. 367–385.

57. Spitz, F. and E.E. Furlong, Transcription factors: from enhancer binding to developmental control. Nat Rev Genet, 2012. 13(9): p. 613–26.

58. Balsalobre, A. and J. Drouin, Pioneer factors as master regulators of the epigenome and cell fate. Nat Rev Mol Cell Biol, 2022. 23(7): p. 449–464.

59. Huang, S., et al., Bifurcation dynamics in lineage-commitment in bipotent progenitor cells. Dev Biol, 2007. 305(2): p. 695–713.

60. Wu, X., Z. Sun, and R. Jiang, Logic motif of combinatorial control in transcriptional networks. Nature Precedings, 2008.

61. Xu, R., et al., Stage-specific H3K9me3 occupancy ensures retrotransposon silencing in human pre-implantation embryos. Cell Stem Cell, 2022. 29(7): p. 1051–1066 e8.

62. Yu, H., et al., Dynamic reprogramming of H3K9me3 at hominoid-specific retrotransposons during human preimplantation development. Cell Stem Cell, 2022. 29(7): p. 1031–1050.e12.

63. Simunovic, M., E.D. Siggia, and A.H. Brivanlou, In vitro attachment and symmetry breaking of a human embryo model assembled from primed embryonic stem cells. Cell Stem Cell, 2022. 29(6): p. 962–972 e4.

64. Pei, W., et al., Resolving Fates and Single-Cell Transcriptomes of Hematopoietic Stem Cell Clones by PolyloxExpress Barcoding. Cell Stem Cell, 2020. 27(3): p. 383–395.e8.

65. Psaila, B., et al., Single-Cell Analyses Reveal Megakaryocyte-Biased Hematopoiesis in Myelofibrosis and Identify Mutant Clone-Specific Targets. Mol Cell, 2020. 78(3): p. 477–492 e8.

66. Aguade-Gorgorio, G., S. Kauffman, and R. Sole, Transition Therapy: Tackling the Ecology of Tumor Phenotypic Plasticity. Bull Math Biol, 2021. 84(1): p. 24.

67. Yang, D., et al., Lineage tracing reveals the phylodynamics, plasticity, and paths of tumor evolution. Cell, 2022. 185(11): p. 1905–1923 e25.

68. Uthamacumaran, A., A review of dynamical systems approaches for the detection of chaotic attractors in cancer networks. Patterns, 2021. 2(4).

69. Tang, Y., et al., TBX20 Improves Contractility and Mitochondrial Function During Direct Human Cardiac Reprogramming. Circulation, 2022: p. 101161CIRCULATIONAHA122059713.

70. Liang, J., et al., Induction of Sertoli-like cells from human fibroblasts by NR5A1 and GATA4. Elife, 2019. 8.

71. Yu, L., et al., Blastocyst-like structures generated from human pluripotent stem cells. Nature, 2021. 591(7851): p. 620-626.

72. Dixit, A., et al., Perturb-Seq: Dissecting Molecular Circuits with Scalable Single-Cell RNA Profiling of Pooled Genetic Screens. Cell, 2016. 167(7): p. 1853–1866 e17.

73. Joung, J., et al., A transcription factor atlas of directed differentiation. Cell, 2023. 186(1): p. 209–229.e26.

74. Replogle, J.M., et al., Mapping information-rich genotype-phenotype landscapes with genome-scale Perturb-seq. Cell, 2022.

75. Domcke, S. and J. Shendure, A reference cell tree will serve science better than a reference cell atlas. Cell, 2023. 186(6): p. 1103–1114.

76. MacArthur, B.D., Stem cell biology needs a theory. Stem Cell Reports, 2023. 18(1): p. 3–5.

77. Casey, M.J., P.S. Stumpf, and B.D. MacArthur, Theory of cell fate. Wiley Interdiscip Rev Syst Biol Med, 2020. 12(2): p. e1471.

78. Stadler, T., O.G. Pybus, and M.P.H. Stumpf, Phylodynamics for cell biologists. Science, 2021. 371(6526).

79. Macnair, W., et al., *Tree*LJ*ensemble analysis assesses presence of multifurcations in single cell data*. Molecular Systems Biology, 2019. 15(3).

80. Shea, M.A. and G.K. Ackers, The OR control system of bacteriophage lambda: A physical-chemical model for gene regulation. Journal of Molecular Biology, 1985. 181(2): p. 211–230.

81. Olariu, V. and C. Peterson, Kinetic models of hematopoietic differentiation. Wiley Interdiscip Rev Syst Biol Med, 2019. 11(1): p. e1424.

82. Yin, J., et al., Construction of a Pathway Map on a Complicated Energy Landscape. Phys Rev Lett, 2020. 124(9): p. 090601.

83. Yin, J., L. Zhang, and P. Zhang, Solution landscape of the Onsager model identifies non-axisymmetric critical points. Physica D: Nonlinear Phenomena, 2022. 430: p. 133081.

84. Yin, J., B. Yu, and L. Zhang, Searching the solution landscape by generalized high-index saddle dynamics. Science China Mathematics, 2020.

85. Mojtahedi, M., et al., Cell Fate Decision as High-Dimensional Critical State Transition. PLoS Biol, 2016. 14(12): p. e2000640.

86. Zhao, Y., Chemically induced cell fate reprogramming and the acquisition of plasticity in somatic cells. Current Opinion in Chemical Biology, 2019. 51: p. 146–153.

87. Brand, M. and E. Morrissey, Single-cell fate decisions of bipotential hematopoietic progenitors. Curr Opin Hematol, 2020. 27(4): p. 232–240.

88. Zhang, Y., et al., Hematopoietic Hierarchy - An Updated Roadmap. Trends Cell Biol, 2018. 28(12): p. 976–986.

89. Arinobu, Y., et al., Reciprocal activation of GATA-1 *and PU.1 marks initial specification of hematopoietic stem cells into myeloerythroid and myelolymphoid lineages.* Cell Stem Cell, 2007. 1(4): p. 416-427.

90. Vanden-Eijnden, E. and M. Heymann, The geometric minimum action method for computing minimum energy paths. The Journal of Chemical Physics, 2008. 128(6): p. 061103.

91. Chen, Q., et al., Tracing the origin of heterogeneity and symmetry breaking in the early mammalian embryo. Nat Commun, 2018. 9(1): p. 1819.

92. de Haan, G. and S.S. Lazare, Aging of hematopoietic stem cells. Blood, 2018. 131(5): p. 479–487.

93. Reya, T., et al., Stem cells, cancer, and cancer stem cells. Nature, 2001. 414(6859): p. 105-111.

94. Shi, Y., et al., Induced pluripotent stem cell technology: a decade of progress. Nat Rev Drug Discov, 2017. 16(2): p. 115–130.

95. Semrau, S., et al., Dynamics of lineage commitment revealed by single-cell transcriptomics of differentiating embryonic stem cells. Nat Commun, 2017. 8(1): p. 1096.

96. Hamazaki, T., et al., Concise Review: Induced Pluripotent Stem Cell Research in the Era of Precision Medicine. Stem Cells, 2017. 35(3): p. 545–550.

97. Abdallah, H.M. and D. Del Vecchio, Computational Analysis of Altering Cell Fate, in Computational Stem Cell Biology: Methods and Protocols, P. Cahan, Editor. 2019, Springer New York: New York, NY. p. 363-405.

98. Qin, J., et al., Direct chemical reprogramming of human cord blood erythroblasts to induced megakaryocytes that produce platelets. Cell Stem Cell, 2022. 29(8): p. 1229–1245.e7.

99. Kumar, L. and M. E Futschik, Mfuzz: a software package for soft clustering of microarray data. Bioinformation, 2007. 2(1): p. 5–7.

100. Cahan, P., et al., Computational Stem Cell Biology: Open Questions and Guiding Principles. Cell Stem Cell, 2021. 28(1): p. 20–32.

101. Del Sol, A. and S. Jung, The Importance of Computational Modeling in Stem Cell Research. Trends Biotechnol, 2021. 39(2): p. 126–136.

102. Lopez-Otin, C., et al., The hallmarks of aging. Cell, 2013. 153(6): p. 1194–217.

103. Beerman, I. and D.J. Rossi, Epigenetic Control of Stem Cell Potential during Homeostasis, Aging, and Disease. Cell Stem Cell, 2015. 16(6): p. 613–25.

104. Dorshkind, K., et al., Do haematopoietic stem cells age? Nat Rev Immunol, 2020. 20(3): p. 196–202.

105. Kovtonyuk, L.V., et al., *Inflamm-Aging of Hematopoiesis,* Hematopoietic Stem Cells, and the Bone Marrow Microenvironment. Front Immunol, 2016. 7: p. 502.

106. Pang, W.W., S.L. Schrier, and I.L. Weissman, Age-associated changes in human hematopoietic stem cells. Semin Hematol, 2017. 54(1): p. 39–42.

107. Wang, P. and L. Chen, Critical transitions and tipping points in EMT. Quantitative Biology, 2020. 8(3): p. 195–202.

108. Scheffer, M., et al., Anticipating Critical Transitions. Science, 2012. 338(6105): p. 344.

109. Liu, R., et al., Identifying critical transitions and their leading biomolecular networks in complex diseases. Sci Rep, 2012. 2: p. 813.

110. Rosales-Alvarez, R.E., et al., Gene expression noise dynamics unveil functional heterogeneity of ageing hematopoietic stem cells. bioRxiv, 2022.

111. Grün, D., L. Kester, and A. van Oudenaarden, Validation of noise models for single-cell transcriptomics. Nat Methods, 2014. 11(6): p. 637–640.

112. Shu, J., et al., Induction of pluripotency in mouse somatic cells with lineage specifiers. Cell, 2013. 153(5): p. 963–75.

113. Saez, M., et al., Statistically derived geometrical landscapes capture principles of decision-making dynamics during cell fate transitions. Cell Syst, 2022. 13(1): p. 12–28 e3.

114. Corson, F. and E.D. Siggia, Geometry, epistasis, and developmental patterning. Proc Natl Acad Sci U S A, 2012. 109(15): p. 5568–75.

115. Fanti, A.-K., et al., Flt3- and Tie2-Cre tracing identifies regeneration in sepsis from multipotent progenitors but not hematopoietic stem cells. Cell Stem Cell, 2023. 30(2): p. 207–218.e7.

