## Supplemental Figure for "A Logic-incorporated Gene Regulatory Network Deciphers Principles in Cell Fate Decisions"

**A Logic-incorporated Gene Regulatory Network**

**Deciphers Principles in Cell Fate Decisions**

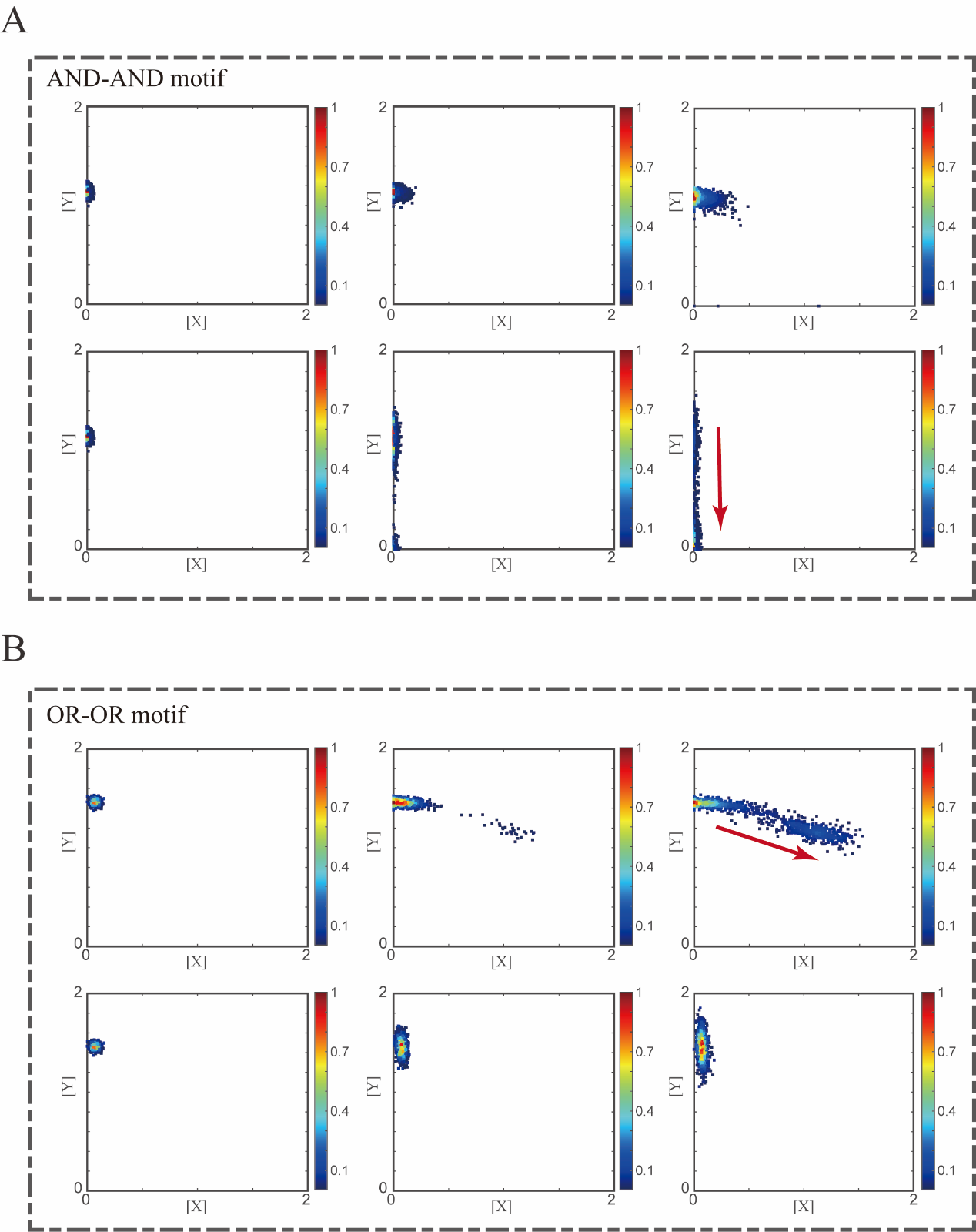

**Figure S1. Noise as a driving force for reprogramming of LY to S in two logic motifs.**

(A) Stochastic simulation in the AND-AND motif. Initial values were identical with attractor of LY fate in Figure 2C top panel (SSS in purple attractor basin). Simulation was preformed 1000 times, with each final state recorded as a dot on the plot. Top panel: Noise level of *X* (*σ_x_*) is set to 0.03, 0.09, 0.15, from left to right, and *σ_y_* is 0.03. Bottom panel: Noise level of *Y* (*σ_y_*) is set to 0.03, 0.09, 0.15, from left to right, and *σ_x_* is 0.03. Red arrow represents the direction of fate transitions of LY to S. Other than adding a white noise, parameters were identical with those in Figure 2C top panel. Color of heatmap corresponds to the density of points.
(B) Stochastic simulation in the OR-OR motifs. Initial values were identical with attractor of LY fate in Figure 2C bottom panel (SSS in blue attractor basin). Simulation was preformed 1000 times, with each final states recorded as a dot on the plot. Top panel: Noise level of *X* (*σ_x_*) is set to 0.03, 0.09, 0.15, from left to right, and *σ_y_* is 0.03. Bottom panel: Noise level of *Y* (*σ_y_*) is set to 0.03, 0.09, 0.15, from left to right, and *σ_x_* is 0.03. Red arrow represents the direction of fate transitions of LY to S. Other than adding a white noise, parameters were identical with these in Figure 2C bottom panel.

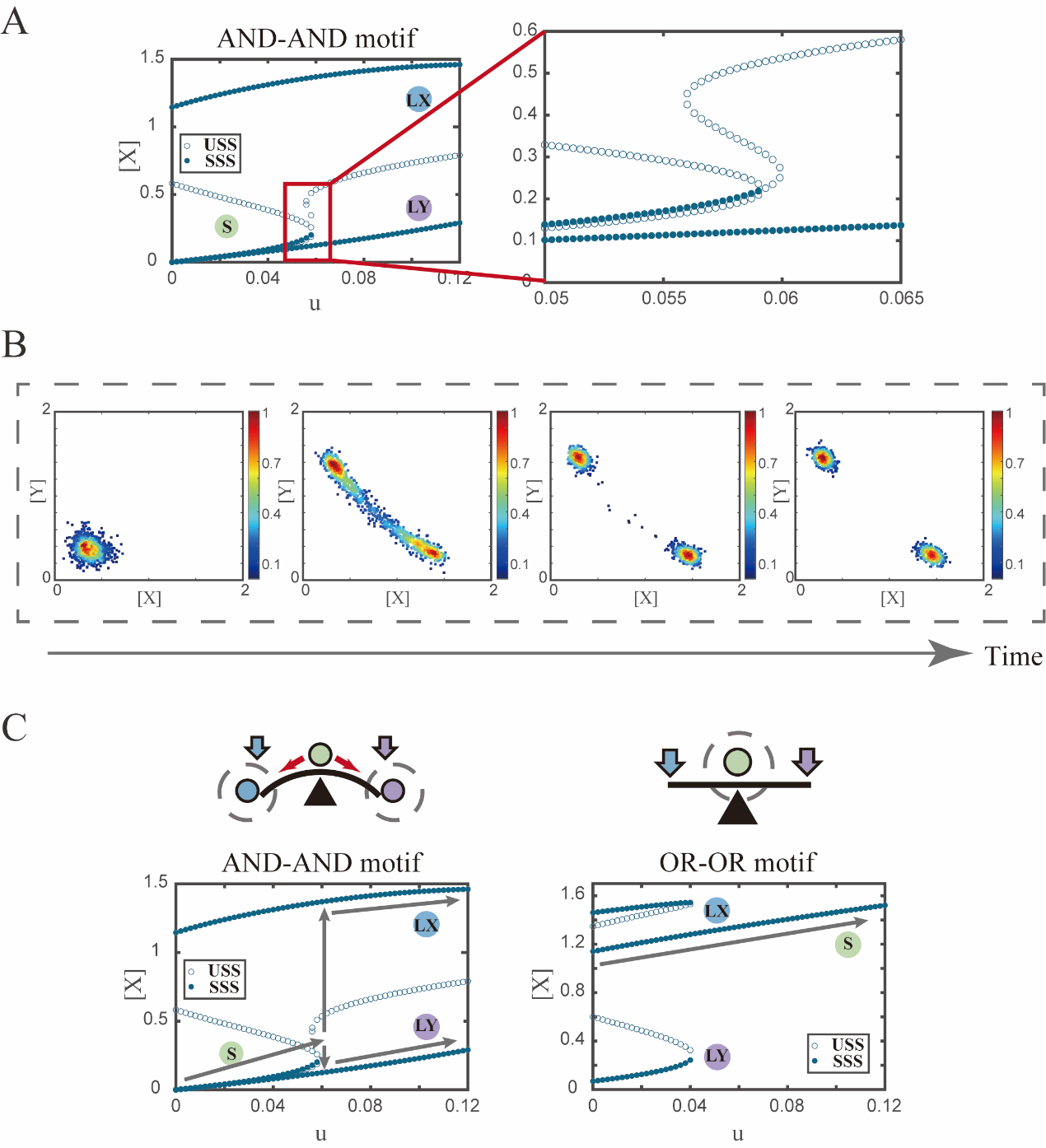

**Figure S2. Two logic motifs are associated with opposite fate-decision choices under bidirectional induction.**

(A) Bifurcation diagrams for the AND-AND motif (**Fig**4.A) driven by parameter *u* (*u* = *u_x_* = *u_y_*) in the CIS model. SSSs and USSs are denoted as solid dots and hollow dots, respectively.

(B) Stochastic simulation in the AND-AND motif. Initial values were identical with attractor of S fate in Figure 2C top panel (SSS in green attractor basin). Noise level of *X* (*σ_x_*) and *Y* (*σ_y_*) are both set to 0.06. Simulation was preformed 1000 times for each pseudo-time point, with each temporal state (from left to right) recorded as a dot on the plot. Model’s parameters were identical with those in Figure 4C 4^th^ panel.

(C) Schematic illustration of the “seesaw” model [1] in two logic motifs under bidirectional induction. Red arrows represent the direction of fate transitions. Blue and purple arrows represent induction of *X* and *Y*, respectively. Grey hollow circles indicate remained cell states. Grey arrows represent fate transitions from S in bifurcation diagrams as increasing *u*.

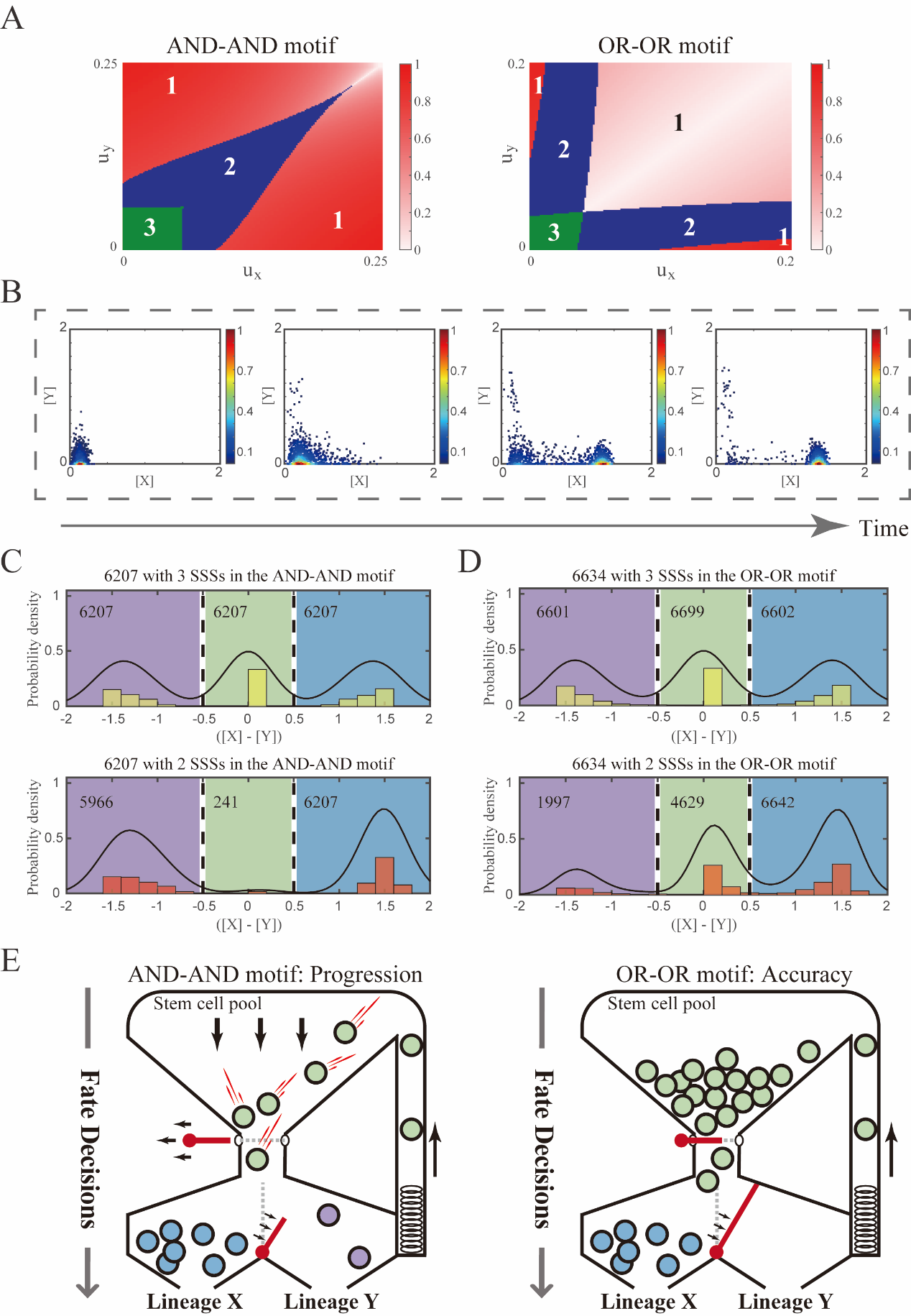

**Figure S3. The PA (progression and accuracy) trade-off of cell fate decisions under the signal-driven mode.**

(A) Phase portraits *u_x_* vs. *u_y_* for the AND-AND and OR-OR motifs. The numbers in separating regions indicate the number of SSS. The extent of red color is quantified by minus of [X] and [Y], which indicates the expression level of balance between *X* and *Y* genes.

(B) Stochastic simulation in the AND-AND motif. Initial values were identical with attractor of S fate in Figure 2C top panel (SSS in green attractor basin). Noise level of *X* (*σ_x_*) and *Y* (*σ_y_*) are set to 0.05, 0.15, respectively. Simulation was preformed 1000 times for each pseudo-time point, with each temporal state (from left to right) recorded as a dot on the plot. Model’s parameters were identical with those in Figure 5D middle panel.

(C-D) Distribution of SSSs under all parameter sets in two logic motifs. We collected parameter sets with 3 SSSs, where 2 SSSs are remained by increasing *u_x_* (see **Methods**). Minus of [X] and [Y] of each SSS is quantified to represent relative cell fate (LX, S, LY). Dark lines are kernel density estimation of each distribution. The numbers in separating regions indicate the number of SSS.

(E) Schematic illustration of PA trade-off. Left panel: the entire red latch of stem cell pool is undone, which indicates all the stem cells are engaged in differentiation with fate bias (the red bar between LX and LY). Right panel: the gate for differentiating into LY is inaccessible, which indicates stem cells “flow” into LX exclusively. Style was inspired by [2] and [3].

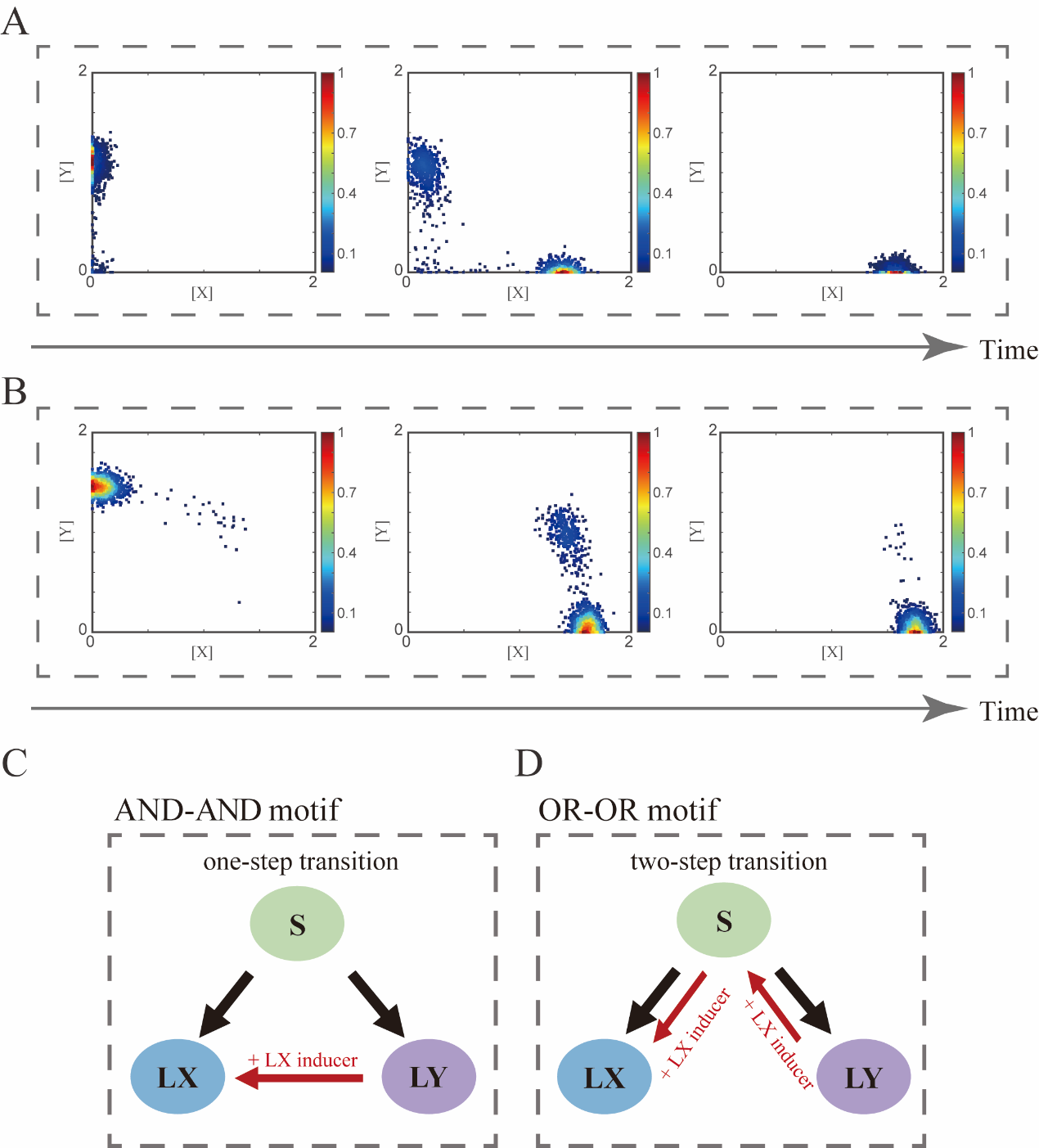

**Figure S4. Distinct trajectories of reprogramming under the signal-driven mode.**

(A-B) Stochastic simulation in the AND-AND (A) and OR-OR (B) motifs. Initial values were identical with attractor of LY fate in Figure 2C (SSSs in purple attractor basins). Noise level of *X* (*σ_x_*) and *Y* (*σ_y_*) are both set to 0.09. Simulation was preformed 1000 times, with each final state recorded as a dot on the plot. Parameter *u_x_* switched from 0 to 0.12 (0, 0.06, 0.12, from left to right). Other model’s parameters were identical with those in Figure 2C.

(C-D) Schematic illustration of trajectories of reprogramming in the AND-AND (C) and OR-OR (D) motifs. Dark arrows indicate in vivo differentiation from stem cell population. Red arrows indicate trajectories of trans-differentiation under the induction of *X*.

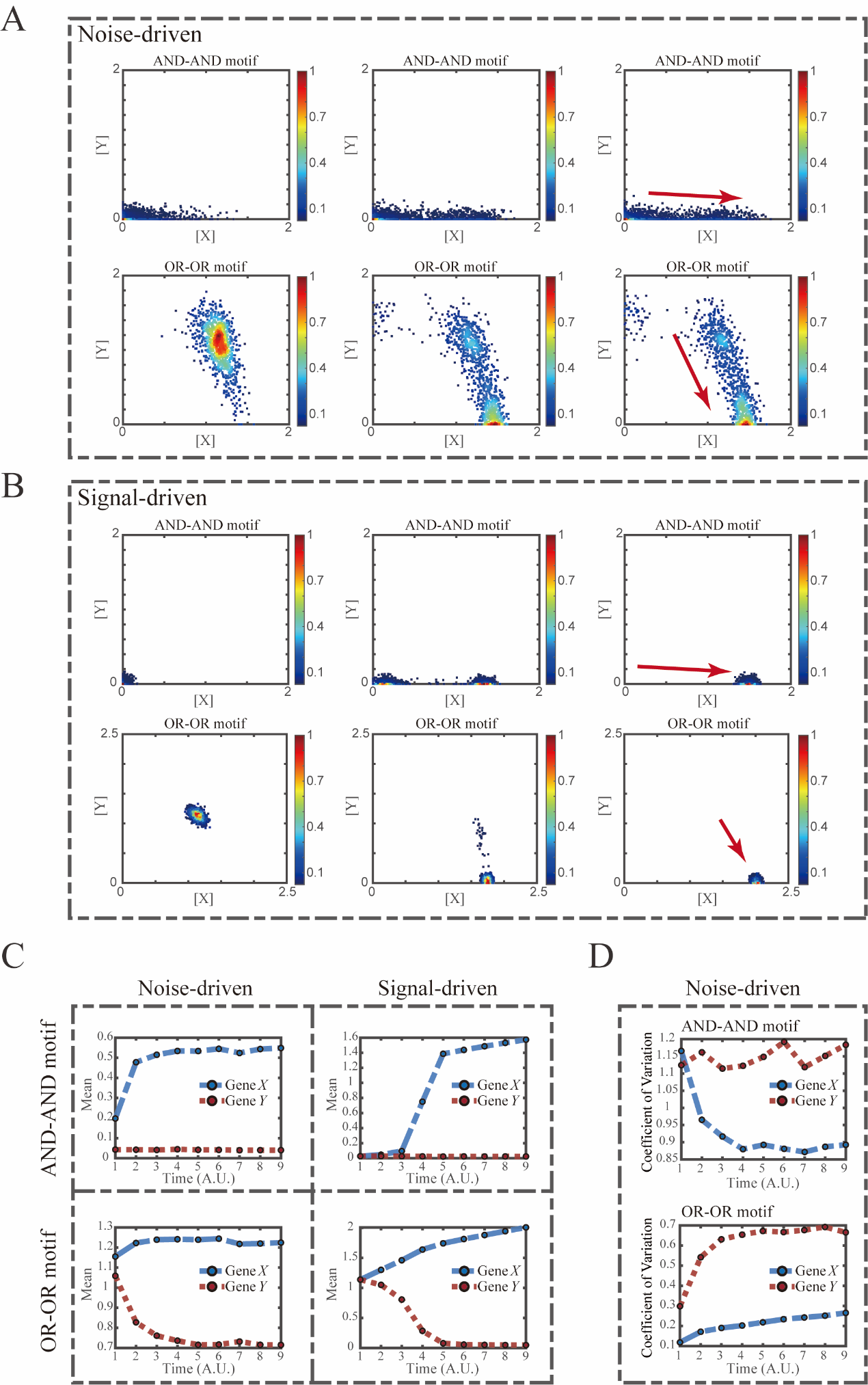

**Figure S5. Simulation of fate transitions of S to LX *in silico* in two logic motifs under two driving forces.**

(A) Simulation in two logic motifs under the noise-driven mode. Initial values were identical with attractor of S fate in Figure 2C (SSSs in green attractor basins). Simulation was preformed 1000 times for each pseudo-time point, with each temporal state (from left to right) recorded as a dot on the plot. Top panel: Noise level of *X* (*σ_x_*) is set to 0.21, and *σ_y_* is 0.09. Bottom panel: Noise level of *Y* (*σ_y_*) is set to 0.21, and *σ_x_* is 0.09. Red arrow represents the direction of fate transitions of S to LX. Other than adding a white noise, parameters were identical with those in Figure 2C.

(B) Simulation in two logic motifs under the signal-driven mode. Initial values were identical with attractor of S fate in Figure 2C (SSSs in green attractor basins). Top panel: Noise level of *X* (*σ_x_*) and *Y* (*σ_y_*) are both set to 0.06. Simulation was preformed 1000 times, with each final state recorded as a dot on the plot. Parameter *u_x_* switched from 0 to 0.09 (0, 0.045, 0.09, from left to right). Bottom panel: Noise level of *X* (*σ_x_*) and *Y* (*σ_y_*) are both set to 0.05. Simulation was preformed 1000 times, with each final state recorded as a dot on the plot. Parameter *u_x_* switched from 0 to 0.24 (0, 0.12, 0.24, from left to right). Red arrow represents the direction of fate transitions of S to LX. Other model’s parameters were identical with those in Figure 2C.

(C) Time courses on the mean in expression levels of *X* and *Y* genes *in silico* during differentiation towards LX. Initial values were set to the attractors of S fate in Figure 2C (SSSs in green attractor basins). Simulation of the noise-driven mode in two logic motifs was identical with that in Figure S5A. Simulation of the signal-driven mode in two logic motifs was identical with that in Figure S5B (Parameter *u_x_* switched from 0 to 0.12 from time point 1 to 9 in the AND-AND motif).

(D) Time courses on the coefficient of variation in expression levels of *X* and *Y* genes *in silico* during differentiation towards LX under the noise-driven mode. Initial values were set to the attractors of S fate in Figure 2C (SSSs in green attractor basins). Simulation of the noise-driven mode in two logic motifs was identical with that in Figure S5A.

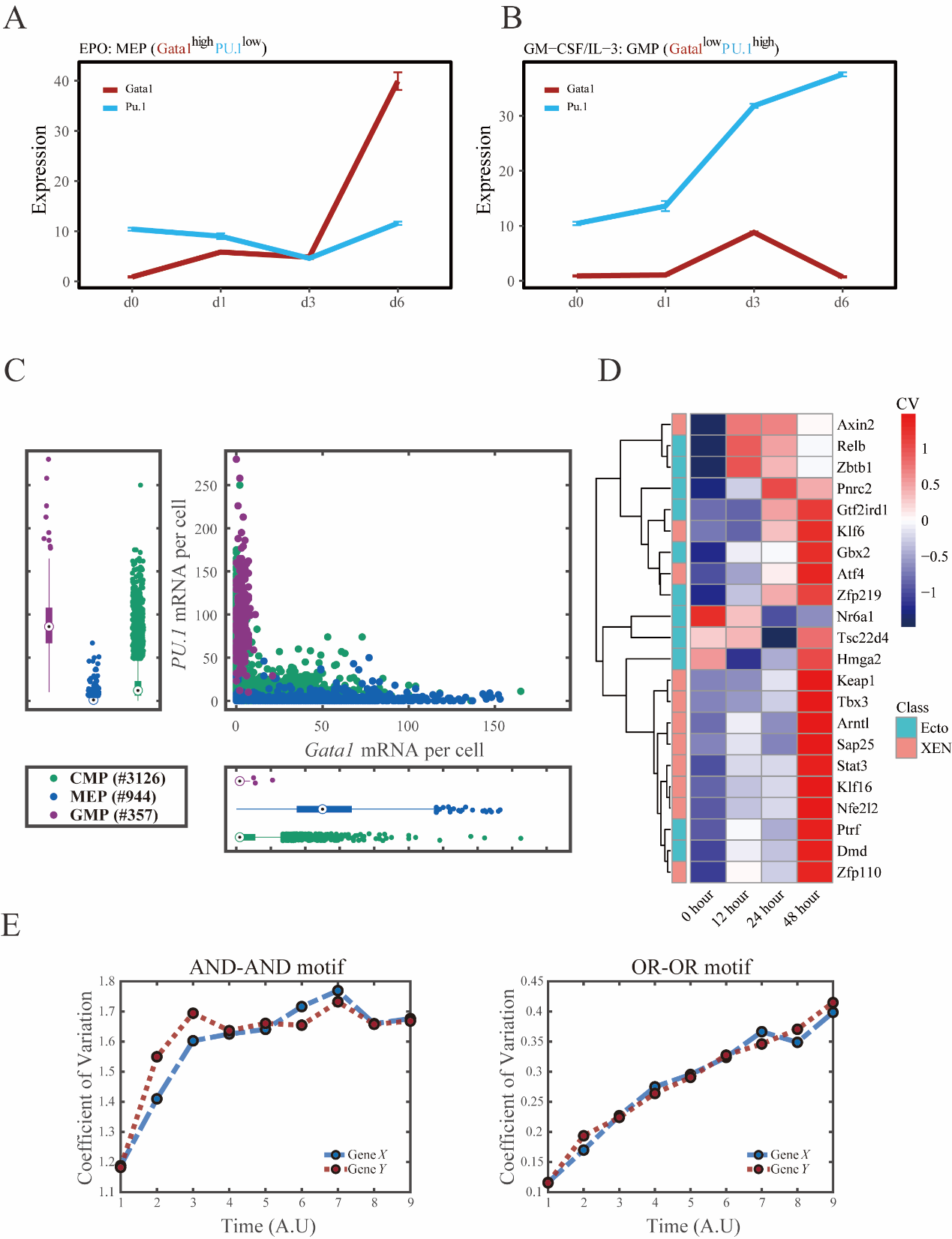

**Figure S6. GRNs perform differently in differentiations during hematopoiesis and embryogenesis.**

(A-B) Measured mean of expression levels of *Gata1* and *PU.1* changing over time in the processes of differentiation from CMPs to MEPs and GMPs. Expression levels were quantified via single-cell RT-qPCR [4]. For details of data processing, see **Methods.**

(C) Expression level of *Gata1* and *PU.1* among CMPs, MEPs and GMPs quantified via single-molecule RNA fluorescent in situ hybridization [5].

(D) Heatmap of coefficient of variation of expression levels of 22 genes among cells in embryogenesis quantified via single-cell SMART-seq2 [6]. For details of data processing, see **Methods.**

(E) Time courses on the coefficient of variation in expression levels of *X* and *Y* genes *in silico* during differentiation under the noise-driven mode. Initial values were set to the attractors of S fate in Figure 2C (SSSs in green attractor basins). Top panel: Noise level of *X* (*σ_x_*) and *Y* (*σ_y_*) are both set to 0.14. Bottom panel: Noise level of *X* (*σ_x_*) and *Y* (*σ_y_*) are both set to 0.1. Stochastic simulation was preformed 1000 times for each pseudo-time point.

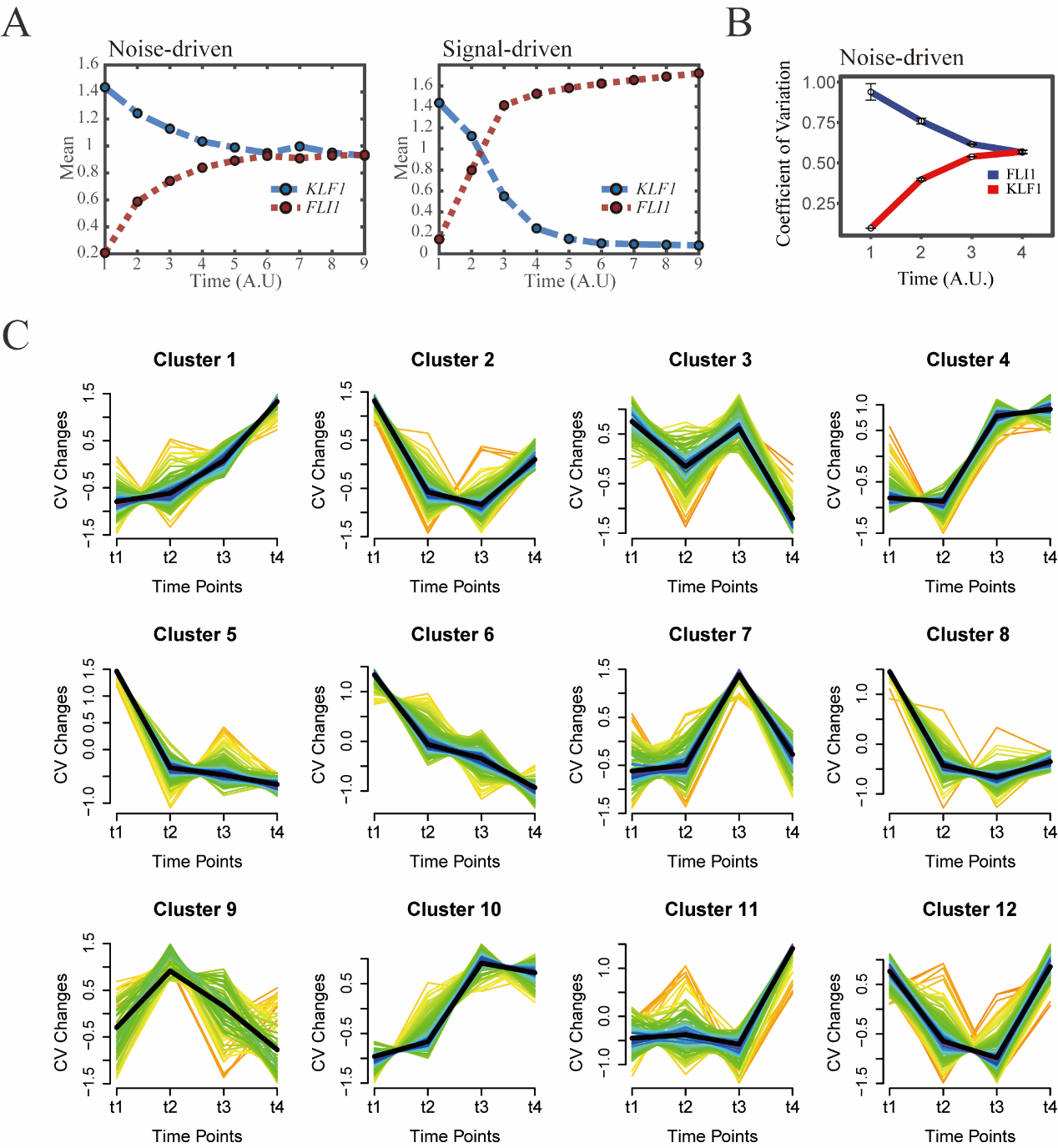

**Figure S7. The chemical-induced reprogramming of human EB to iMK is the signal-driven fate decisions with an OR-OR-like motif.**

(A) Time courses on the mean in expression levels of *KLF1* and *FLI1* genes *in silico* during reprogramming of EBs to iMKs under the noise-driven (left) and signal-driven (right) modes. Initial values were set to the attractors of LX fate in Figure 2C bottom panel (SSS in blue attractor basin). Noise level of *KLF1* (*σ_x_*) and *FLI1* (*σ_y_*) are both set to 0.18 under the noise-driven mode. Stochastic simulation was preformed 1000 times for each pseudo-time point. Other than adding a white noise, parameters were identical with those in Figure 2C bottom panel. Simulation of the signal-driven mode was identical with that in Figure 7E left panel.

(B) Coefficient of variation of expression levels of *KLF1* and *FLI1* changes *in silico* along by pseudo-time under the OR-OR motif. Simulation was identical with that in Figure S7A left panel.

(C) Identification of distinct temporal patterns of expression variance by fuzzy c-means clustering. The x axis represents four time points, while the y axis represents scaled CV (coefficient of variation) in each time point. Dark trend lines in the middle indicate average of scaled CV over genes in cluster.

**Reference**

1.	Shu, J., et al., *Induction of pluripotency in mouse somatic cells with lineage specifiers.* Cell, 2013. **153**(5): p. 963-75.

2.	Chen, Q., et al., *Tracing the origin of heterogeneity and symmetry breaking in the early mammalian embryo.* Nat Commun, 2018. **9**(1): p. 1819.

3.	Goldberg, A.D., C.D. Allis, and E. Bernstein, *Epigenetics: a landscape takes shape.* Cell, 2007. **128**(4): p. 635-8.

4.	Mojtahedi, M., et al., *Cell Fate Decision as High-Dimensional Critical State Transition.* PLoS Biol, 2016. **14**(12): p. e2000640.

5.	Wheat, J.C., et al., *Single-molecule imaging of transcription dynamics in somatic stem cells.* Nature, 2020. **583**(7816): p. 431-436.

6.	Semrau, S., et al., *Dynamics of lineage commitment revealed by single-cell transcriptomics of differentiating embryonic stem cells.* Nat Commun, 2017. **8**(1): p. 1096.
