## Supplemental Method for "A Logic-incorporated Gene Regulatory Network Deciphers Principles in Cell Fate Decisions"

#### Contents


**1 Derivation of the CIS network** **1**

**2 Stochastic simulations** **5**

**3 Parameter screening** **6**

**4 Saddle points and saddle dynamics** **6**

**5 Constructing the solution landscape** **7**

**6 Screening the fully-connected stage** **7**

**7 Data processing of single-cell data** **8**

**8 Fuzzy C-means Clustering** **9**

**9 GO enrichment analysis** **9**

**10 Integration of human TF-target interactions** **9**

**11 Integration of human TF list** **9**


#### 1 Derivation of the CIS network

In GRN, each TFs is represented by a node and the edge between nodes represent the regulatory relationship. In our work, we considered a GRN comprised of 2 TFs (i.e., X, Y), formulated with 2 ODEs describing the change of each TF. Define ($[X]$,$[Y]$) to be the concentration of TF X,Y, the ODE model is described as follows:

$$\begin{cases} \dfrac{d[X]}{dt} = H_X([X],[Y]) - d_X[X] \\ \dfrac{d[Y]}{dt} = H_Y([X],[Y]) - d_Y[Y], \end{cases} \tag{1}$$

where $H_X([X],[Y])$ is the production rate of TF $X$ that combines the effects from both activators and suppressors of the X and $d_X$ is the decay rate from the $X$ and $Y$, which integrate degradation and dilution.

The exact form of $H_X([X],[Y]), H_Y([X],[Y])$ are derived based on the simple GRN with nodes $X$ and $Y$ via a set of molecular interactions between these TFs themselves, genes that encode for them, and the mRNAs[1].

TFs in a GRN act as multimers to implement regulatory interaction. In our model, we treated TFs X and Y as acting in their homo-multimer forms, $X_{n_1}$ and $Y_{n_2}$, respectively (i.e., $n_1$ TF X monomers reversibly form a activated homomultizerized form $X_{n_1}$ and $n_2$ monomers of TF Y to reversibly form activated homo-multimer $Y_{n_2}$). Of note, $n_1$ and $n_2$ here are able to be further generalized beyond the number of binding elements[2, 3]. The multimerization biochemical reaction of TFs X and Y can be represented as follows,

$$\begin{aligned} \underbrace{X + X + \cdots + X}_{\text{n}_1 \text{ monomers}} &\underset{q_x}{\overset{p_x}{\rightleftharpoons}} X_{n_1}, \\ \underbrace{Y + Y + \cdots + Y}_{\text{n}_2 \text{ monomers}} &\underset{q_y}{\overset{p_y}{\rightleftharpoons}} Y_{n_2}. \end{aligned} \tag{2}$$

There are many diverse mechanisms of regulation[4, 5]. For simplicity, our model considered transcriptional regulation as major mechanism since it is feasibly characterized in experiments and can well represent the interaction between genes in **C**ross-**I**nhibition with **S**elf activation (CIS) network. To activate downstream transcription, TFs bind with their Cis-Regulatory Elements (CREs). We denoted as $D_X$ for the no-bound CREs of TF X and $D_Y$ for the TF Y in an independent manner. Here we posited different TFs bind to exclusive, non-overlapping CREs to regulate target genes. Hence, there are totally eight binding patterns described as follows,

$$\begin{aligned} D_X + X_{n_1} &\underset{q_1}{\overset{p_1}{\rightleftharpoons}} D_{XX}, & D_X + Y_{n_2} &\underset{q_2}{\overset{p_2}{\rightleftharpoons}} D_{XY}, \\ D_Y + Y_{n_2} &\underset{q_4}{\overset{p_4}{\rightleftharpoons}} D_{YY}, & D_Y + X_{n_1} &\underset{q_3}{\overset{p_3}{\rightleftharpoons}} D_{YX}, \\ D_{XX} + Y_{n_2} &\underset{q_5}{\overset{p_5}{\rightleftharpoons}} D_{XXY}, & D_{XY} + X_{n_1} &\underset{q_6}{\overset{p_6}{\rightleftharpoons}} D_{XXY}, \\ D_{YX} + Y_{n_2} &\underset{q_7}{\overset{p_7}{\rightleftharpoons}} D_{YYX}, & D_{YY} + X_{n_1} &\underset{q_8}{\overset{p_8}{\rightleftharpoons}} D_{YYX}. \end{aligned} \tag{3}$$

Next, we modeled the biochemical reactions of transcription and translation. Transcription is an elaborate process involving many steps, including initiation, elongation, and termination of an mRNA transcript. Likewise, translation includes peptide formation and elongation, followed by protein folding before function. However, to evaluate the behavior of GRN concisely, we treated transcription and translation as single-step reactions and then use lumped rates to encompass the time it takes for all steps in the elaborate machinery to complete[1]. Hence, the transcription process can be represented as follows,

$$\begin{aligned} & & D_{XX} &\xrightarrow{\alpha_x^1} m_X + D_{XX}, & D_{YY} &\xrightarrow{\alpha_y^1} m_Y + D_{YY}, \\ D_X &\xrightarrow{\alpha_x} m_X + D_X, & D_{XY} &\xrightarrow{\alpha_x^2} m_X + D_{XY}, & D_{YX} &\xrightarrow{\alpha_y^2} m_Y + D_{YX}, \\ D_Y &\xrightarrow{\alpha_y} m_Y + D_Y, & D_{XXY} &\xrightarrow{\alpha_x^3} m_X + D_{XXY}, & D_{YYX} &\xrightarrow{\alpha_y^3} m_Y + D_{YYX}. \end{aligned} \tag{4}$$

Here, we distinguished the basal rate of transcription of a gene without activation or repression (termed as constitutive transcription). The basal transcription rate is shown in the left panel in 4.

Though we considered eight configurations, these ultimately led to two mRNA transcripts, $m_X$ and $m_Y$. Translation can also be represented by a one-step process from these transcripts to their protein products. The one-step translation biochemical reactions are described as follows,

$$m_X \xrightarrow{\kappa_x} m_X + X, \quad m_Y \xrightarrow{\kappa_y} m_Y + Y. \tag{5}$$

Next, we considered the decay of these proteins as well as their respective mRNA species. In general, these proteins and mRNA species will undergo decay to some extent which is a combination of both degradation and dilution.

$$\begin{aligned} X &\xrightarrow{\delta_x} \emptyset, & m_X &\xrightarrow{\eta_x} \emptyset, & X &\xrightarrow{\beta} \emptyset, \\ Y &\xrightarrow{\delta_y} \emptyset, & m_Y &\xrightarrow{\eta_y} \emptyset, & Y &\xrightarrow{\beta} \emptyset, \end{aligned} \tag{6}$$

here, the left and middle panels represent the degradation of TFs X and Y as well as mRNA transcripts $m_X$ and $m_Y$. The right panel represents the dilution of TFs X and Y due to cell division. In general, the degradation rate $\gamma$ can be determined using the following relations,

$$\delta = \frac{\ln 2}{t_{1/2}}, \quad \eta = \frac{\ln 2}{t_{1/2}}, \quad \beta \approx \frac{\ln 2}{t_{doubling}}, \tag{7}$$

where $t_{1/2}$ represents the half-lives of each protein or mRNA transcript, $t_{doubling}$ represents the cell division rate. For protein products X and Y, we used the notation $d$ to illustrate the total rate of decay rather than 2 separate parameters in our model, i.e., $d_1 = \delta_x + \beta$, $d_2 = \delta_y + \beta$, respectively. In general, $t_{1/2}$ and $t_{doubling}$ are robust and stationary, thus we treated these paramaters as constant in our model, i.e., $d_i = \text{Const}, \text{ for } i = 1, 2$.

We have described the reactions that comprise the endogenous components of our GRN. In our work, fate decisions are classified into two modes, i.e., driven by the noise of intracellular gene expression or driven by extracellular signals. Under the noise-driven mode, a gaussian white noise is added to the concentration of TFs to illustrate the random intracellular noise. Whereas if driven by signals, the expression of a gene will be affected by extracellular molecular cocktails, physical stimulation, etc. Therefore, we added the production rate of the TF's mRNA from the ectopic DNA to generalize our model.

$$\emptyset \overset{u_x}{\to} m_X, \emptyset \overset{u_y}{\to} m_Y, \tag{8}$$

here $u_x$, $u_y$ represent the additional mRNA species $m_X$ and $m_Y$ via ectopic overexpression at rates $u_x$ and $u_y$, respectively.

The biochemical reactions take place on a faster timescale. This enables us to derive the change in concentration of species from a biochemical reaction based on the law of mass action. For example, for multimerization reaction of X in Eq2, once equilibrium, we can get

$$\underbrace{X + X + \cdots + X}_{\text{n}_1 \text{ monomers}} \underset{q_x}{\overset{p_x}{\rightleftharpoons}} X_{n_1} \Rightarrow p_x \underbrace{[X][X]\cdots[X]}_{\text{n}_1 \text{ terms}} = p_x[X]^{n_1} = q_x[X_{n_1}] \Rightarrow K_x \triangleq \frac{p_x}{q_x} = \frac{[X_{n_1}]}{[X]^{n_1}}. \tag{9}$$

Also, for TF X bind with its CRE $D_X$, we can get

$$D_X + X_{n_1} \underset{q_1}{\overset{p_1}{\rightleftharpoons}} D_{XX} \Rightarrow p_1[D_X][X_{n_1}] = q_1[D_{XX}] \Rightarrow K_1 \triangleq \frac{p_1}{q_1} = \frac{[D_{XX}]}{[D_X][X_{n_1}]}, \tag{10}$$

here we define $K_1 \triangleq \dfrac{p_1}{q_1}$ as a constant, i.e., chemical equilibrium constant. Furthermore, we define $K_i \triangleq \dfrac{p_i}{q_i},\ i = x, y, 1, 2, \cdots, 8$ for biochemical reactions in Eq2 and Eq3, respectively.

Then, we constructed ODE model for the GRN. For a given species $S$, the principle to get a single equation is as follows:

$$\text{Change in concentration of } S = \frac{d[S]}{dt} = [\dot{S}] = \sum\{\text{all biochemical reaction rates involving } S\}. \tag{11}$$

To explain how this principle assists us to build the ODE model, we took species $m_X$ as a running example. Summarizing all of these biochemical reaction rates involving $m_X$ we can get the change in concentration $[m_X]$,

$$[\dot{m_X}] = \alpha_x[D_X] + \alpha_x^1[D_{XX}] + \alpha_x^2[D_{XY}] + \alpha_x^3[D_{XXY}] - \eta_x[m_X] + u_x. \tag{12}$$

Doing so for our biochemical reaction network yielded the equations of our ODE model,

$$\begin{cases} \dot{[X]} = \kappa_x[m_X] - d_1[X] \\ \dot{[Y]} = \kappa_y[m_Y] - d_2[Y] \\ \dot{[m_X]} = \alpha_x[D_X] + \alpha_x^1[D_{XX}] + \alpha_x^2[D_{XY}] + \alpha_x^3[D_{XXY}] - \eta_x[m_X] + u_x \\ \dot{[m_Y]} = \alpha_y[D_Y] + \alpha_y^1[D_{YY}] + \alpha_y^2[D_{YX}] + \alpha_y^3[D_{YYX}] - \eta_y[m_Y] + u_y \\ \dot{[D_X]} = -p_1[X_{n_1}][D_X] + q_1[D_{XX}] - p_2[Y_{n_2}][D_X] + q_2[D_{XY}] \\ \dot{[D_Y]} = -p_4[Y_{n_2}][D_Y] + q_4[D_{YY}] - p_3[X_{n_1}][D_Y] + q_3[D_{YX}] \\ \dot{[D_{XX}]} = p_1[D_X][X_{n_1}] - q_1[D_{XX}] - p_5[D_{XX}][Y_{n_2}] + q_5[D_{XXY}] \\ \dot{[D_{XY}]} = p_2[D_X][Y_{n_2}] - q_2[D_{XY}] - p_6[D_{XY}][X_{n_1}] + q_6[D_{XXY}] \\ \dot{[D_{YX}]} = p_3[D_Y][X_{n_1}] - q_3[D_{YX}] - p_7[D_{YX}][Y_{n_2}] + q_7[D_{YYX}] \\ \dot{[D_{YY}]} = p_4[D_Y][Y_{n_2}] - q_4[D_{YY}] - p_8[D_{YY}][X_{n_1}] + q_8[D_{YYX}] \\ \dot{[D_{XXY}]} = p_5[D_{XX}][Y_{n_2}] - q_5[D_{XXY}] + p_6[D_{XY}][X_{n_1}] - q_6[D_{XXY}] \\ \dot{[D_{YYX}]} = p_7[D_{YX}][Y_{n_2}] - q_7[D_{YYX}] + p_8[D_{YY}][X_{n_1}] - q_8[D_{YYX}] \\ \dot{[X_{n_1}]} = p_x[X]^{n_1} - q_x[X_{n_1}] - p_1[D_X][X_{n_1}] + q_1[D_{XX}] - p_3[D_Y][X_{n_1}] + q_3[D_{YX}] \\ \qquad - p_6[D_{XY}][X_{n_1}] + q_6[D_{XXY}] - p_8[D_{YY}][X_{n_1}] + q_8[D_{YYX}] \\ \dot{[Y_{n_2}]} = p_y[Y]^{n_2} - q_y[Y_{n_2}] - p_2[D_X][Y_{n_2}] + q_2[D_{XY}] - p_4[D_Y][Y_{n_2}] + q_4[D_{YY}] \\ \qquad - p_5[D_{XX}][Y_{n_2}] + q_5[D_{XXY}] - p_7[D_{YX}][Y_{n_2}] + q_7[D_{YYX}]. \end{cases} \tag{13}$$

The 14-dimension ODE model of the GRN contains overwhelming parameters that make it impractical to analyze for cell fate. To reduce the dimensionality of the model, we assumed the multimerization, DNA binding/unbinding and mRNA dynamic occurs sufficiently faster than protein production and decay, the temporal derivatives of the respective species can be set to 0, indicating that the species concentration reaches its quasi-steady state[1]. Thus, we can get

$$\begin{cases} \dot{[X]} = \kappa_x[m_X] - d_1[X] \\ \dot{[Y]} = \kappa_y[m_Y] - d_2[Y] \\ \dot{[m_X]} = 0 = \alpha_x[D_X] + \alpha_x^1[D_{XX}] + \alpha_x^2[D_{XY}] + \alpha_x^3[D_{XXY}] - \eta_x[m_X] + u_x \\ \dot{[m_Y]} = 0 = \alpha_y[D_Y] + \alpha_y^1[D_{YY}] + \alpha_y^2[D_{YX}] + \alpha_y^3[D_{YYX}] - \eta_y[m_Y] + u_y. \end{cases} \tag{14}$$

To better demonstrate the deviation of the model, we take TF X as a running example. Since GRN is symmetric, we can deviate TF Y's equation in the same way. Once the biochemical reaction reaches its equilibrium, using the law of mass action we can get

$$\begin{aligned} [X_{n_1}] &= K_x[X]^{n_1} \\ [D_{XX}] &= K_1[D_X][X_{n_1}] = K_1[D_X][X_{n_1}] = K_xK_1[D_X][X]^{n_1}, \\ [D_{XY}] &= K_2[D_X][Y_{n_2}] = K_2[D_X][Y_{n_2}] = K_yK_2[D_X][Y]^{n_2}, \\ [D_{XXY}] &= K_5[D_{XX}][Y_{n_2}] = K_5[D_{XX}][Y_{n_2}] = K_xK_yK_1K_5[D_X][X]^{n_1}[Y]^{n_2}. \end{aligned} \tag{15}$$

Noticing that $\dot{[m_X]} = 0$ in Eq14, the species $[m_X]$ can be represented by CRE terms,

$$\begin{aligned} [m_X] &= \frac{1}{\eta_x}\left(\alpha_x[D_X] + \alpha_x^1[D_{XX}] + \alpha_x^2[D_{XY}] + \alpha_x^3[D_{XXY}]\right) + \frac{u_x}{\eta_x} \\ &= \frac{1}{\eta_x}\left(\alpha_x[D_X] + \alpha_x^1K_1[D_X][X_{n_1}] + \alpha_x^2K_2[D_X][Y_{n_2}] + \alpha_x^3K_1K_5[D_X][X_{n_1}][Y_{n_2}]\right) + \frac{u_x}{\eta_x} \\ &= \frac{1}{\eta_x}[D_X]\left(\alpha_x + \alpha_x^1K_xK_1[X]^{n_1} + \alpha_x^2K_yK_2[Y]^{n_2} + \alpha_x^3K_xK_yK_1K_5[X]^{n_1}[Y]^{n_2}\right) + \frac{u_x}{\eta_x}. \end{aligned} \tag{16}$$

In our well-stirred system, for TF X's CREs, the conservation law holds, i.e., the total CRE concentration of X, as a constant, equals the sum of CRE concentration that is bound by X, by

Y, and by X and Y. By employing so and combining with Eq15 we can get a representation of $[D_X]$,

$$[D_{TX}] = [D_X]+[D_{XY}]+[D_{XX}]+[D_{XXY}] \Rightarrow [D_X] = \frac{[D_{TX}]}{(1+K_xK_1[X]^{n_1}+K_yK_2[Y]^{n_2}+K_xK_yK_1K_5[X]^{n_1}[Y]^{n_2})}. \tag{17}$$

Substitute $[D_X]$ in Eq16 with Eq17, we can get

$$\begin{aligned} [m_X] &= \frac{1}{\eta_x}[D_X]\left(\alpha_x+\alpha_x^1K_xK_1[X]^{n_1}+\alpha_x^2K_yK_2[Y]^{n_2}+\alpha_x^3K_xK_yK_1K_5[X]^{n_1}[Y]^{n_2}\right)+\frac{u_x}{\eta_x} \\ &= \frac{D_{TX}}{\eta_x}\frac{\left(\alpha_x+\alpha_x^1K_xK_1[X]^{n_1}+\alpha_x^2K_yK_2[Y]^{n_2}+\alpha_x^3K_xK_yK_1K_5[X]^{n_1}[Y]^{n_2}\right)}{(1+K_xK_1[X]^{n_1}+K_yK_2[Y]^{n_2}+K_xK_yK_1K_5[X]^{n_1}[Y]^{n_2})}+\frac{u_x}{\eta_x}. \end{aligned} \tag{18}$$

Then finally we reach to the point that the deviation of $[\dot{X}]$. In Eq14 we substituted $[m_X]$ with Eq18, and got

$$\begin{aligned} [\dot{X}] &= \kappa_x[m_X]-d_1[X] \\ &= \frac{D_{TX}\kappa_x}{\eta_x}\frac{\left(\alpha_x+\alpha_x^1K_xK_1[X]^{n_1}+\alpha_x^2K_yK_2[Y]^{n_2}+\alpha_x^3K_xK_yK_1K_5[X]^{n_1}[Y]^{n_2}\right)}{(1+K_xK_1[X]^{n_1}+K_yK_2[Y]^{n_2}+K_xK_yK_1K_5[X]^{n_1}[Y]^{n_2})}+\frac{\kappa_xu_x}{\eta_x}-d_1[X], \end{aligned} \tag{19}$$

here, we define $r_0^x = \alpha_x\frac{D_{TX}\kappa_x}{\eta_x}, r_1 = \alpha_x^1\frac{D_{TX}\kappa_x}{\eta_x}, r_2 = \alpha_x^2\frac{D_{TX}\kappa_x}{\eta_x}, r_3 = \alpha_x^3\frac{D_{TX}\kappa_x}{\eta_x}$ as relative protein generation rates, $k_1 = K_1K_x, k_2 = K_yK_2, k_3 = K_xK_yK_1K_5$ as the weight of each TF binding patterns, $\tilde{u}_x = \frac{\kappa_xu_x}{\eta_x}$ as ectopic DNA (to simplify the notation, we may drop the tilde of $\widetilde{u_i},\ i=x,y$), then we can get

$$[\dot{X}] = \frac{r_0^x+r_1k_1[X]^{n_1}+r_2k_2[Y]^{n_2}+r_3k_3[X]^{n_1}[Y]^{n_2}}{1+k_1[X]^{n_1}+k_2[Y]^{n_2}+k_3[X]^{n_1}[Y]^{n_2}}-d_1[X]+u_x, \tag{20}$$

Similarly, for TF Y, we can get

$$\begin{aligned} [\dot{Y}] &= \frac{D_{TY}\kappa_y}{\eta_y}\frac{\left(\alpha_y+\alpha_y^1K_yK_3[Y]^{n_2}+\alpha_y^2K_yK_4[X]^{n_1}+\alpha_y^3K_xK_yK_3K_7[X]^{n_1}[Y]^{n_2}\right)}{(1+K_yK_3[Y]^{n_2}+K_yK_4[X]^{n_1}+K_xK_yK_3K_7[X]^{n_1}[Y]^{n_2})}+\frac{\kappa_yu_y}{\eta_y}-d_2[Y] \\ &= \frac{r_0^y+r_4k_4[Y]^{n_2}+r_5k_5[X]^{n_1}+r_6k_6[X]^{n_1}[Y]^{n_2}}{1+k_4[Y]^{n_2}+k_5[X]^{n_1}+k_6[X]^{n_1}[Y]^{n_2}}-d_2[Y]+u_y. \end{aligned} \tag{21}$$

Ultimately, combining Eq20 and Eq21, we reduced our 14-dimension model to a 2-dimension dynamic system,

$$\begin{cases} \frac{d[X]}{dt} = \frac{r_0^x+r_1k_1[X]^{n_1}+r_2k_2[Y]^{n_2}+r_3k_3[X]^{n_1}[Y]^{n_2}}{1+k_1[X]^{n_1}+k_2[Y]^{n_2}+k_3[X]^{n_1}[Y]^{n_2}}-d_1[X]+u_x \\ \frac{d[Y]}{dt} = \frac{r_0^y+r_4k_4[Y]^{n_2}+r_5k_5[X]^{n_1}+r_6k_6[X]^{n_1}[Y]^{n_2}}{1+k_4[Y]^{n_2}+k_5[X]^{n_1}+k_6[X]^{n_1}[Y]^{n_2}}-d_2[Y]+u_y, \end{cases} \tag{22}$$

### 2 Stochastic simulations

In Eq22, parameters of the AND-AND motif (Fig2.C top panel) are: $(r_0^x, r_0^y, r_1, r_2, r_3, r_4, r_5, r_6)$=(0, 0, 0.95, 0, 0, 0.95, 0, 0), $(k_1, k_2, k_3, k_4, k_5, k_6) = (1.5, 0.8, 1, 1.5, 0.8, 1)$ and $(d_1, d_2) = (0.55, 0.55)$. Parameters of the OR-OR motif (Fig2.C bottom panel) are: $(r_0^x, r_0^y, r_1, r_2, r_3, r_4, r_5, r_6)$=(0.15, 0.15, 0.95, 0, 0.95, 0.95, 0, 0.95), $(k_1, k_2, k_3, k_4, k_5, k_6) = (2.1, 1.7, 1.8, 2.1, 1.7, 1.8)$ and $(d_1, d_2) = (0.55, 0.55)$. In both the AND-AND and OR-OR motifs, $(n_1, n_2) = (2, 2)$. We simulated noise in the model using a Langevin equation[6]:

$$dx_i = F_i(\mathbf{x})dt + dW_i, \tag{23}$$

Where $F_i(\mathbf{x})$ is the deterministic function (22). $W_i$ is a Wiener process which introduces additive noise with no dependence on the state $\mathbf{x}$. We integrated the Wiener process over the interval which gives us $\sqrt{\Delta t}\xi_i$, and $\xi_i$ is a Gaussian white noise with zero mean and given variance $(\sigma^2)$. Thus we can computed the new state of the dynamical systems using the Runge–Kutta method.

### 3 Parameter screening

We conducted global parameters screening of the fully-connected stage (FCS) and Progression-Accuracy trade-off in our models. First, to collect parameter sets with 3 SSSs, we used Latin hypercube sampling (LHS) to screen k-series parameters symmetrically (i.e., $k_1 = k_4, k_2 = k_5, k_3 = k_6$) ranging from 0.001 to 5 both in the AND-AND and OR-OR motifs. We ultimately collected 6,231 sets for the AND-AND motif and 6,682 sets for the OR-OR motifs (Table S1).

To analyze the sequence of vanishing SSSs. We further filtered parameter sets with 2 SSSs remained as increasing $u_x$ (6,207 sets for the AND-AND motif; 6,634 sets for the OR-OR motif). For each SSS, we quantified the Minus of [X] and [Y] (FigS3.C-D).

### 4 Saddle points and saddle dynamics

Set an autonomous dynamical system[7]

$$\dot{\boldsymbol{x}} = \boldsymbol{F}(\boldsymbol{x}), \boldsymbol{x} \in R^n, \tag{24}$$

where $\boldsymbol{F} : \mathbb{R}^n \to \mathbb{R}^n$ is a $\mathcal{C}^r (r \geqslant 2)$ function, and a point $\boldsymbol{x}^* \in \mathbb{R}^n$is called a stationary point (or equilibrium solution) of Eq24 if $\boldsymbol{F}(\boldsymbol{x}^*) = 0$. Let $J(\boldsymbol{x}) = \nabla \boldsymbol{F}(\boldsymbol{x})$ denote the Jacobian of $\boldsymbol{F}(\boldsymbol{x})$. For a stationary point $\boldsymbol{x}^*$, taking $\boldsymbol{x} = \boldsymbol{x}^* + \boldsymbol{y}$ in Eq24, we have

$$\dot{\boldsymbol{y}} = J\left(\boldsymbol{x}^*\right) \boldsymbol{y} + \mathcal{O}\left(\|\boldsymbol{y}\|^2\right) \tag{25}$$

where $\| * \|$ denotes the norm induced by the inner product. The associated linear system

$$\dot{\boldsymbol{y}} = J\left(\boldsymbol{x}^*\right) \boldsymbol{y}, \tag{26}$$

is used to determine the stability of $\boldsymbol{x}^*$. Depending on the eigenvalues of $J(\boldsymbol{x})$ with positive, negative and zero real parts, we can define unstable, stable and center manifolds of the Jacobian $J(\boldsymbol{x})$ spanned by the corresponding eigenvectors, as $\mathcal{W}^{\mathrm{u}}\left(\boldsymbol{x}^*\right), \mathcal{W}^{\mathrm{s}}\left(\boldsymbol{x}^*\right)$ and $\mathcal{W}^{\mathrm{c}}\left(\boldsymbol{x}^*\right)$.

From the primary decomposition theorem, $\mathbb{R}^n$can be decomposed as a direct sum:

$$\mathbb{R}^n = \mathcal{W}^{\mathrm{u}}(\boldsymbol{x}) \oplus \mathcal{W}^{\mathrm{s}}(\boldsymbol{x}) \oplus \mathcal{W}^{\mathrm{c}}(\boldsymbol{x}), \tag{27}$$

A hyperbolic stationary point is called a saddle if $\mathcal{W}^{\mathrm{u}}\left(\boldsymbol{x}^*\right)$ and $\mathcal{W}^{\mathrm{s}}\left(\boldsymbol{x}^*\right)$ are nontrivial. The hyperbolic stationary point $\boldsymbol{x}^*$ is called a sink (source) if all the eigenvalues of $J\left(\boldsymbol{x}^*\right)$ have negative (positive) real parts. The index of a stationary point $\boldsymbol{x}^*$ is defined as the dimension of the unstable subspace $\mathcal{W}^{\mathrm{u}}\left(\boldsymbol{x}^*\right)$

The HiOSD method[8] is designed for finding index-k saddles of an energy function $E(\boldsymbol{x})$. For gradient systems, $\boldsymbol{F}(\boldsymbol{x}) = -\nabla E(\boldsymbol{x})$, and the Jacobian $J(\boldsymbol{x}) = -\nabla^2 E(\boldsymbol{x}) = -G(\boldsymbol{x})$, where $G(\boldsymbol{x})$ denotes the Hessian of $E(\boldsymbol{x})$. The k-saddle $\boldsymbol{x}^*$ is a local maximum on the linear manifold $\boldsymbol{x}^*+ \mathcal{W}^{\mathrm{u}}\left(\boldsymbol{x}^*\right)$ and a local minimum on $\boldsymbol{x}^* + \mathcal{W}^{\mathrm{s}}\left(\boldsymbol{x}^*\right)$. So, the high-index saddle dynamics (HiSD) for a index k saddle( k-saddle) is a transformed gradient flow

$$\dot{\boldsymbol{x}} = -\mathcal{P}_{\mathcal{W}^{\mathrm{u}}(\boldsymbol{x})} \boldsymbol{F}(\boldsymbol{x}) + \left(\boldsymbol{F}(\boldsymbol{x}) - \mathcal{P}_{\mathcal{W}^{\mathrm{u}}(\boldsymbol{x})} \boldsymbol{F}(\boldsymbol{x})\right) = \left(I - 2\mathcal{P}_{\mathcal{W}^{\mathrm{u}}(\boldsymbol{x})}\right) \boldsymbol{F}(\boldsymbol{x}), \tag{28}$$

where $\mathcal{P}_{\mathcal{V}}$ denotes the orthogonal projection operator on a finite-dimensional subspace $\mathcal{V}$. Here, $-\mathcal{P}_{\mathcal{W}^{\mathrm{u}}(\boldsymbol{x})} \boldsymbol{F}(\boldsymbol{x})$ is taken as an ascent direction on the subspace $\mathcal{W}^{\mathrm{u}}(\boldsymbol{x})$ and $\boldsymbol{F}(\boldsymbol{x}) - \mathcal{P}_{\mathcal{W}^{\mathrm{u}}(\boldsymbol{x})} \boldsymbol{F}(\boldsymbol{x})$ is a descent direction on the subspace $\mathcal{W}^{\mathrm{S}}(\boldsymbol{x})$. The subspace $\mathcal{W}^{\mathrm{u}}(\boldsymbol{x}) = \operatorname{span}\{\boldsymbol{v_1}, \ldots, \boldsymbol{v_k}\}$ where $\boldsymbol{v_i}$ is the unit eigenvector corresponding to the smallest $\boldsymbol{i}$-th eigenvalues, which can be obtained by many methods such as minimize the Rayleigh quotients. According to the above the HiSD for a k-saddle (k-HiSD) is:

$$\left\{\begin{array}{l} \beta^{-1} \dot{\boldsymbol{x}} = \left(\mathbb{I} - \sum_{i=1}^k 2 \boldsymbol{v}_i \boldsymbol{v}_i^{\top}\right) \boldsymbol{F}(\boldsymbol{x}) \\ \gamma^{-1} \dot{\boldsymbol{v}}_i = -\left(\mathbb{I} - \boldsymbol{v}_i \boldsymbol{v}_i^{\top} - \sum_{j=1}^{i-1} 2 \boldsymbol{v}_j \boldsymbol{v}_j^{\top}\right) \mathbb{G}(\boldsymbol{x}) \boldsymbol{v}_i, i = 1, 2, \ldots, k, \end{array}\right. \tag{29}$$

which coupled with the initial condition

$$\boldsymbol{x}(0) = \boldsymbol{x}^{(0)} \in \mathbb{R}^n, \boldsymbol{v}_i(0) = \boldsymbol{v}_i^{(0)} \in \mathbb{R}^n, \text{ s.t. } \left\langle \boldsymbol{v}_j^{(0)}, \boldsymbol{v}_i^{(0)} \right\rangle = \delta_{ij}, i, j = 1, \ldots, k, \tag{30}$$

Where $\mathbb{I}$ is the identity operator and $\beta, \gamma > 0$ are relaxation parameters.

Similarly, the GHiSD[7] for a k-saddle (k-GHiSD) of the dynamical system 24 has the following form:

$$\begin{cases} \dot{\boldsymbol{x}} = \left(I - 2\sum_{j=1}^{k} \boldsymbol{v}_j \boldsymbol{v}_j^{\mathrm{T}}\right) \boldsymbol{F}(\boldsymbol{x}) \\ \dot{\boldsymbol{v}}_i = \left(I - \boldsymbol{v}_i \boldsymbol{v}_i^{\mathrm{T}}\right) J(\boldsymbol{x})\boldsymbol{v}_i - \sum_{j=1}^{i-1} \boldsymbol{v}_j \boldsymbol{v}_j^{\mathrm{T}} \left(J(\boldsymbol{x}) + J^{\mathrm{T}}(\boldsymbol{x})\right) \boldsymbol{v}_i, i = 1, \dots, k, \end{cases} \tag{31}$$

which coupled with the initial condition 30. The k-GHiSD can be accelerated by Heavy Ball method.

### 5 Constructing the solution landscape

For a given set of parameters {a1...,u}, we can use k-GHiSD to find each order saddle point of the dynamical system, and then construct the solution landscape of the system. The solution landscape is a pathway map consisting of all stationary points and their connections[9, 10]. The solution landscape is a new tool used to describe the dynamic behavior of stationary points in a dynamic system which can show the connection and transfer path between stationary points. In general, we can use the downward search algorithm and the upward search algorithm to construct the solution landscape, which starting from the k-saddle points and the stable points respectively. Because of the symmetry and other prior knowledge of the dynamic system used in our paper, The saddle points of the system tend to occur in the range of $[0, x_{max}]^2$, where the $x_{max}$ means the maximum coordinate of the stable point. We grid the range, select each grid point as the initial state, and find saddle points of corresponding order ($k = 1, 2$) using k-GHiSD. Taking the saddle points found before as the initial states, we disturb its unstable directions to search other stationary points of lower index, establish the connection relationship between stationary points and finally construct the solution landscape. In addition, we use geometric minimum action method (gMAM)[11] to find the minimum action path between the stable points, use the obtained action to represent the stability of the stable points, and represent them at different heights in the solution landscape.

### 6 Screening the fully-connected stage

As shown in the case of $u = 0.0565$ in Fig4.D, the fully-connected stage has three stable points and three 1-index saddle points to connect the stable points with each other. Therefore, we posited that finding the fully-connected stage can be equivalent to finding three different 1-index saddle points in this system.

Firstly, build a set of parameters with three stable points. In two different logic motifs, 10,000 sets of dynamic parameters are randomly generated respectively, and all stable points of the system corresponding to each set of parameters are found. If there are three stable points, the corresponding dynamic parameters are recorded as a new set of parameters for the subsequent search of 1-index saddle points.

Secondly, finding the 1-index saddle points in a system with three stable points. According to the above results, we can find that saddle points tend to appear in the range of $[0, x_{max}]^2$, where the $x_{max}$ means the maximum coordinate of the stable point. So we can grid the range in the same process as before, and search 1-index saddle points with each grid point as the initial structure. In order to find the relevant saddle points as detailed as possible, denser grid points were used as the initial structure of the system without three 1-index saddle points to find the 1-index saddle points.

Due to the calculation error of numerical calculation itself, the saddle points found by saddle point dynamics may have the following problems:

1. The same saddle points found from different initial structures were mistaken for two.
2. Finding a negative or infinite saddle point.
3. The saddle-point judgment is inaccurate due to the influence of zero eigenvalue.

We can set corresponding judgment conditions to solve the above problems.

In addition, since there is also a parameter $u$ representing the foreign signal in the system, we can let $u$ vary to find if there are three 1-index saddle points. At the beginning, $u = 0$ is set, and then $u$ is changed according to the number of saddle points found and the corresponding judgment conditions, so as to find the system with three 1-index saddle points. We use the idea of adaptive step size to speed up the search process in the algorithm of finding three 1-index saddle points in the system. The specific process is as follows:

- Set the initial $u = 0$, $du = 1$.
- Saddle Dynamics is used to find the 1-index saddle points in the parameters.
- When the number of 1-index saddle points is equal to 0: there is only one stable point in the system ($u = 0.12$ in Fig4.F), it indicates that $u$ is too large and needs to be reduced to make $u = u - du$.
- When the number of 1-index saddle points is equal to 1: the system has two stable points ($u = 0.12$ in Fig4.C), it indicates that $u$ is too large and needs to be reduced so that $u = u - du$.
- When the number of 1-index saddle points is equal to 2: the system has two kinds of cases, one of which needs to make $u$ increase ($u = 0$ in Fig4.C). In this case, the coordinate of stable point near $y = x$ is smaller than the midpoint of two 1-index saddle points, so we let $u = u + du$, and let $du = 0.1 * du$. In the other case, there are two possibilities. One is like $u = 0$ in Fig4.F, and the other is like $u = 0.0595$ in Fig4.F. In this case, there is no stable point near $y = x$ or the coordinate of the stable point is larger than the midpoint of two 1-index saddle points. It indicates that $u$ is too large and needs to be reduced so that $u = u - du$
- When the system finds that the number of 1-index saddle points is equal to 3: we need to judge whether there are three non-degenerate saddle points that are different from each other. If the condition is sufficient, it indicates that we have found the fully-connected solution landscape; otherwise, let $u = u + du$, $du = 0.1 * du$, and continue searching.
- When the number of 1-index saddle points is greater than 3, it indicates that $u$ is too large and needs to be reduced to make $u = u - du$.
- If three 1-index saddle points are not found after 100 changes of $u$, it is considered that there is no fully-connected solution landscape in the system.

We respectively used the above algorithm to search for the fully-connected solution landscape under two different logic motifs and obtained the following results:

| Logic | # of 3 SSS | # of FCS | Ratio |
|---|---|---|---|
| AND-AND | 6231 | 5151 | 82.7% |
| OR-OR | 6682 | 120 | 1.8% |

It can be seen from the simulation results that the fully-connected solution landscape can be captured in most AND-AND motifs, but almost not in the OR-OR motifs. Due to the error of numerical calculation itself, the obtained results may have a certain deviation. In addition, in our search process, due to the limitation of computing resources and the complex nature of some systems, in order to take into account the majority of cases, some systems with full connectivity have not been found. Subsequent manual search found that the fully-connected solution landscape exists in systems which not found under the AND-AND motifs previously (17.3%). Therefore, the obtained proportion can be further improved.

### 7 Data processing of single-cell data

There are totally 4 public datasets used in our work (related to Fig6.F and H, Fig7, FigS6.C). In mice's hematopoiesis[12], expression of genes is quantified as $2^{LOD-Cq}-1$ (LOD: limit of dection; Cq-value is the cutoff of amplification signal in single-cell qPCR data). Thus we can computed the coefficient of variation (CV) over time. In embryogenesis[13], public dataset used in our work was generated by single-cell SMART-seq2 (GEO: GSE79578). We utilized preprocessed data to compute the CV over time. In reprogramming[14], public dataset was generated by 10x Genomics (GEO: GSE207654). Counts Per Million (CPM) matrix are used to compute the CV. We assigned time points by Leiden clustering algorithm with parameters consistent with original paper (see methods in [14]).

### 8 Fuzzy C-means Clustering

CV of 2,000 high variable genes given by original paper[14] over four time points were computed. We filtered out 1,677 genes by removing NA due to missing values. Noise of 1,677 genes were then grouped into 12 clusters using Mfuzz package in R with fuzzy c-means algorithm[15] (Table S2).

### 9 GO enrichment analysis

GO and pathway enrichment analyses were performed using Metascape[16].

### 10 Integration of human TF-target interactions

A collection of 456,698 TF-target interactions for 1,541 human TFs and their mode of regulation (MoR, activation or repression) were integrated by merging a built-in collection of human regulons from the DoRothEA R package (version 1.8.0)[17] and human TRRUST database[18]. To be specific, resource from DoRothEA includes 454,504 TF-target interactions for 1,541 human TFs, retrieved from (1) literature-curated resources, (2) ChIP-seq binding data, (3) TFBS (TF binding sites) predictions, and (4) transcriptional regulatory interactions inferred from published gene expression profiles. Human TRRUST database is a manually curated database of transcriptional regulatory networks, whose current version contains 8,427 TF-target regulatory relationships of 795 human TFs with annotation of MoR (activation, repression or unknown). For our purpose, we excluded TF-target interactions with unknown MoR, yet 207 TF-target pairs still remained showing conflict MoRs in records from literature, according to TRRUST database. To integrate data from both resources, when a conflict occurred in TRRUST database, we retained the MoR recorded in DoRothEA resource for the same pair of TF-target, but omitted the conflict records if the TF-target pair was not included in DoRothEA resource. As for TF-target pairs without conflict MoRs in TRRUST database but shows conflicts across the two resources, records in TRRUST database were taken as more credible (Table S3).

### 11 Integration of human TF list

In addition to the 1,541 human TFs we obtained from the collection of TF-target interactions mentioned earlier, 1,639 human TFs reported by Lambert et al.[5] and 1,564 by Ng et al.[19] were used as complements, ending up with a total of 2,051 human TFs (Table S3).

### References


[1] Hussein M. Abdallah and Domitilla Del Vecchio. *Computational Analysis of Altering Cell Fate*, pages 363–405. Springer New York, New York, NY, 2019.

[2] Kee-Myoung Nam, Rosa Martinez-Corral, and Jeremy Gunawardena. The linear framework: using graph theory to reveal the algebra and thermodynamics of biomolecular systems. *Interface Focus*, 12(4):20220013, 2022.

[3] M. Santillán. On the use of the hill functions in mathematical models of gene regulatory networks. *Mathematical Modelling of Natural Phenomena*, 3(2):85–97, 2008.

[4] A. Balsalobre and J. Drouin. Pioneer factors as master regulators of the epigenome and cell fate. *Nat Rev Mol Cell Biol*, 23(7):449–464, 2022.

[5] S. A. Lambert, A. Jolma, L. F. Campitelli, P. K. Das, Y. Yin, M. Albu, X. Chen, J. Taipale, T. R. Hughes, and M. T. Weirauch. The human transcription factors. *Cell*, 172(4):650–665, 2018.

[6] David V. Foster, Jacob G. Foster, Sui Huang, and Stuart A. Kauffman. A model of sequential branching in hierarchical cell fate determination. *Journal of Theoretical Biology*, 260(4):589–597, 2009.

[7] Jianyuan Yin, Bing Yu, and Lei Zhang. Searching the solution landscape by generalized high-index saddle dynamics. *Science China Mathematics*, 2020.

[8] Jianyuan Yin, Lei Zhang, and Pingwen Zhang. High-index optimization-based shrinking dimer method for finding high-index saddle points. *SIAM Journal on Scientific Computing*, 41(6):A3576–A3595, 2019.

[9] J. Yin, Y. Wang, J. Z. Y. Chen, P. Zhang, and L. Zhang. Construction of a pathway map on a complicated energy landscape. *Phys Rev Lett*, 124(9):090601, 2020.

[10] Jianyuan Yin, Lei Zhang, and Pingwen Zhang. Solution landscape of the onsager model identifies non-axisymmetric critical points. *Physica D: Nonlinear Phenomena*, 430:133081, 2022.

[11] Eric Vanden-Eijnden and Matthias Heymann. The geometric minimum action method for computing minimum energy paths. *The Journal of Chemical Physics*, 128(6):061103, 2008.

[12] M. Mojtahedi, A. Skupin, J. Zhou, I. G. Castano, R. Y. Leong-Quong, H. Chang, K. Trachana, A. Giuliani, and S. Huang. Cell fate decision as high-dimensional critical state transition. *PLoS Biol*, 14(12):e2000640, 2016.

[13] S. Semrau, J. E. Goldmann, M. Soumillon, T. S. Mikkelsen, R. Jaenisch, and A. van Oudenaarden. Dynamics of lineage commitment revealed by single-cell transcriptomics of differentiating embryonic stem cells. *Nat Commun*, 8(1):1096, 2017.

[14] Jinhua Qin, Jian Zhang, Jianan Jiang, Bowen Zhang, Jisheng Li, Xiaosong Lin, Sihan Wang, Meiqi Zhu, Zeng Fan, Yang Lv, Lijuan He, Lin Chen, Wen Yue, Yanhua Li, and Xuetao Pei. Direct chemical reprogramming of human cord blood erythroblasts to induced megakaryocytes that produce platelets. *Cell Stem Cell*, 29(8):1229–1245.e7, 2022.

[15] Lokesh Kumar and Matthias E Futschik. Mfuzz: a software package for soft clustering of microarray data. *Bioinformation*, 2(1):5–7, 2007.

[16] Yingyao Zhou, Bin Zhou, Lars Pache, Max Chang, Alireza Hadj Khodabakhshi, Olga Tanaseichuk, Christopher Benner, and Sumit K. Chanda. Metascape provides a biologist-oriented resource for the analysis of systems-level datasets. *Nature Communications*, 10(1):1523, 2019.

[17] L. Garcia-Alonso, C. H. Holland, M. M. Ibrahim, D. Turei, and J. Saez-Rodriguez. Benchmark and integration of resources for the estimation of human transcription factor activities. *Genome Res*, 29(8):1363–1375, 2019.

[18] Heonjong Han, Jae-Won Cho, Sangyoung Lee, Ayoung Yun, Hyojin Kim, Dasom Bae, Sunmo Yang, Chan Yeong Kim, Muyoung Lee, Eunbeen Kim, Sungho Lee, Byunghee Kang, Dabin Jeong, Yaeji Kim, Hyeon-Nae Jeon, Haein Jung, Sunhwee Nam, Michael Chung, Jong-Hoon Kim, and Insuk Lee. Trrust v2: an expanded reference database of human and mouse transcriptional regulatory interactions. *Nucleic Acids Research*, 46(D1):D380–D386, 2018.

[19] A. H. M. Ng, P. Khoshakhlagh, J. E. Rojo Arias, G. Pasquini, K. Wang, A. Swiersy, S. L. Shipman, E. Appleton, K. Kiaee, R. E. Kohman, A. Vernet, M. Dysart, K. Leeper, W. Saylor, J. Y. Huang, A. Graveline, J. Taipale, D. E. Hill, M. Vidal, J. M. Melero-Martin, V. Busskamp, and G. M. Church. A comprehensive library of human transcription factors for cell fate engineering. *Nat Biotechnol*, 39(4):510–519, 2021.
